# N-terminal domain antigenic mapping reveals a site of vulnerability for SARS-CoV-2

**DOI:** 10.1101/2021.01.14.426475

**Authors:** Matthew McCallum, Anna De Marco, Florian Lempp, M. Alejandra Tortorici, Dora Pinto, Alexandra C. Walls, Martina Beltramello, Alex Chen, Zhuoming Liu, Fabrizia Zatta, Samantha Zepeda, Julia di Iulio, John E. Bowen, Martin Montiel-Ruiz, Jiayi Zhou, Laura E. Rosen, Siro Bianchi, Barbara Guarino, Chiara Silacci Fregni, Rana Abdelnabi, Shi-Yan Caroline Foo, Paul W. Rothlauf, Louis-Marie Bloyet, Fabio Benigni, Elisabetta Cameroni, Johan Neyts, Agostino Riva, Gyorgy Snell, Amalio Telenti, Sean P.J. Whelan, Herbert W. Virgin, Davide Corti, Matteo Samuele Pizzuto, David Veesler

## Abstract

SARS-CoV-2 entry into host cells is orchestrated by the spike (S) glycoprotein that contains an immunodominant receptor-binding domain (RBD) targeted by the largest fraction of neutralizing antibodies (Abs) in COVID-19 patient plasma. Little is known about neutralizing Abs binding to epitopes outside the RBD and their contribution to protection. Here, we describe 41 human monoclonal Abs (mAbs) derived from memory B cells, which recognize the SARS-CoV-2 S N-terminal domain (NTD) and show that a subset of them neutralize SARS-CoV-2 ultrapotently. We define an antigenic map of the SARS-CoV-2 NTD and identify a supersite recognized by all known NTD-specific neutralizing mAbs. These mAbs inhibit cell-to-cell fusion, activate effector functions, and protect Syrian hamsters from SARS-CoV-2 challenge. SARS-CoV-2 variants, including the 501Y.V2 and B.1.1.7 lineages, harbor frequent mutations localized in the NTD supersite suggesting ongoing selective pressure and the importance of NTD-specific neutralizing mAbs to protective immunity.

## Introduction

The emergence of SARS-CoV-2 coronavirus at the end of 2019 resulted in the ongoing COVID-19 pandemic, bringing the world to a standstill (Zhou et al., 2020; Zhu et al., 2020). The lack of pre-existing immunity to SARS-CoV-2 combined with its efficient human-to-human transmission has already resulted in more than 86 million infections and over 1.85 million fatalities as of January 2021. Although vaccines are being developed and deployed at an unprecedented pace, the timeline for large-scale manufacturing and distribution to a large enough population for achieving community protection remain uncertain. As a result, prophylactic and/or therapeutic anti-viral drugs are expected to play a role in controlling COVID-19 disease and the ongoing pandemic. Such drugs may be helpful for unvaccinated individuals or those who respond poorly to vaccination as well as upon waning of immunity or emergence of antigenically distinct strains.

SARS-CoV-2 infects host cells through attachment of the viral transmembrane spike (S) glycoprotein to angiotensin-converting enzyme 2 (ACE2) followed by fusion of the viral and host membranes (Letko et al., 2020; Walls et al., 2020c; Wrapp et al., 2020; Zhou et al., 2020). SARS-CoV-2 S also engages cell-surface heparan-sulfates (Clausen et al., 2020), neuropilin-1 (Cantuti-Castelvetri et al., 2020; Daly et al., 2020) and L-SIGN/DC-SIGN (Chiodo et al., 2020; Gao et al., 2020; Soh et al., 2020; Thépaut et al., 2020) which were proposed to serve as co-receptors, auxiliary receptors, or adsorption factors. SARS-CoV-2 S is the main target of neutralizing Abs in infected individuals and the focus of the many nucleic acid, vectored, and protein subunit vaccines currently deployed or in development (Corbett et al., 2020a; Corbett et al., 2020b; Erasmus et al., 2020; Hassan et al., 2020; Keech et al., 2020; Mercado et al., 2020; Walls et al., 2020b). Besides blocking ACE2 attachment (Piccoli et al., 2020; Tortorici et al., 2020), some neutralizing Abs are alternatively expected to interfere with heparan-sulfate, neuropilin-1 or L-SIGN/DC-SIGN interactions.

The SARS-CoV-2 S protein comprises an N-terminal S_1_ subunit responsible for virus–receptor binding, and a C-terminal S_2_ subunit that promotes virus–cell membrane fusion (Walls et al., 2020c; Wrapp et al., 2020). The S_1_ subunit comprises an N-terminal domain (NTD) and a receptor-binding domain (RBD), also known as domain A and B, respectively (Tortorici and Veesler, 2019). Antibodies targeting the RBD account for 90% of the neutralizing activity in COVID-19 convalescent sera (Piccoli et al., 2020) and numerous monoclonal antibodies (mAbs) recognizing this domain have been isolated and characterized (Barnes et al., 2020a; Barnes et al., 2020b; Baum et al., 2020b; Brouwer et al., 2020; Hansen et al., 2020; Ju et al., 2020; Piccoli et al., 2020; Pinto et al., 2020; Tortorici et al., 2020; Wang et al., 2020; Wu et al., 2020). Several RBD-specific mAbs capable of protecting small animals and non-human primates from SARS-CoV-2 challenge are able to neutralize viral infection by targeting multiple distinct antigenic sites (Baum et al., 2020a; Hansen et al., 2020; Jones et al., 2020; Pinto et al., 2020; Rogers et al., 2020; Tortorici et al., 2020; Zost et al., 2020). A subset of these mAbs is currently being evaluated in clinical trials or have recently received emergency use authorization from the FDA.

The limited immunogenicity of the SARS-CoV-2 NTD in COVID-19 patients (Piccoli et al., 2020; Rogers et al., 2020) has been hypothesized to result from its extensive N-linked glycan shielding (Walls et al., 2020c; Watanabe et al., 2020). However, recent studies have reported on the isolation of NTD-targeted mAbs and their ability to neutralize SARS-CoV-2 infection *in vitro* suggesting they could be useful for COVID-19 prophylaxis or treatment (Chi et al., 2020; Liu et al., 2020a). Although the NTD has been proposed to interact with auxiliary receptors in cell types that do not express ACE2 (e.g. DC-SIGN/L-SIGN), its role and the mechanism of action of NTD targeted neutralizing mAbs remain unknown (Soh et al., 2020). Understanding the immunogenicity of different S domains and the function of mAbs targeting them, including the NTD, is critical to a full understanding of immunity during the pandemic.

Here, we analyze Ab responses in three COVID-19 convalescent individuals and describe 41 NTD-specific human mAbs. Integrating cryo-electron microscopy (cryoEM), binding assays, and antibody escape mutants analysis we define a SARS-CoV-2 NTD antigenic map and identify a supersite recognized by potent neutralizing mAbs. We show that these mAbs exhibit neutralization activities on par with best-in-class RBD-specific mAbs and efficiently activate Fc-mediated effector functions. We also identify immunologically important variation of the SARS-CoV-2 NTD suggesting that the S glycoprotein is under selective pressure from the host humoral immune response. Finally, we provide proof-of-principle that a highly potent NTD mAb provides prophylactic protection against lethal SARS-CoV-2 challenge of Syrian hamsters paving the way for evaluating this class of mAbs in the clinic.

### NTD-specific mAbs with potent neutralizing activity

To discover mAbs targeting diverse SARS-CoV-2 epitopes, we sorted IgG^+^ memory B cells from peripheral blood mononuclear cells (PBMCs) of three COVID-19 convalescent individuals (L, M, X) using biotinylated prefusion SARS-CoV-2 S as a bait. The percentage of SARS-CoV-2 S-reactive IgG^+^ B cells ranged between 1.1 - 1.3 % of IgG^+^ memory B cells. A total of 278 mAbs were isolated and recombinantly produced as human IgG1 (**Figure 1A)**. Characterization by ELISA showed that most mAbs isolated from the three donors recognize the RBD (65-77%), with a smaller fraction targeting the NTD (6-20%). The remaining mAbs (4-20%) are expected to bind to either the S_2_ subunit or the C-D domains within the S_1_ subunit **(Figure 1A)**. The low proportion of NTD-specific mAbs isolated from these donors is in line with the previously observed limited NTD immunogenicity in SARS-CoV-2 exposed individuals (Piccoli et al., 2020; Rogers et al., 2020). Overall, we identified 41 mAbs recognizing the SARS-CoV2 NTD with EC50s ranging between 7.6 - 698 ng/ml and nanomolar binding affinities, as evaluated using ELISA and biolayer interferometry, respectively **(Figure 1B, S1A, Table S1**,**and SI Items 1)**. These NTD-specific mAbs use a large repertoire of V genes, with an over-representation of IGHV3-21 and IGK3-15 genes **(Figure S1B-C and Table S1)**. These mAbs harbor few somatic hypermutations (VH and VL are 97.57% and 97.54% identical to V germline genes, respectively; **Figure S1D, Table S1)**, as previously described for most SARS-CoV-2 neutralizing mAbs binding to the RBD (Piccoli et al., 2020; Seydoux et al., 2020). CDRH3 lengths range between 10 and 24 amino acid residues **(Figure S1E)**. Collectively, these data indicate that the Ab response to the SARS-CoV-2 NTD is polyclonal.

**Figure 1.**
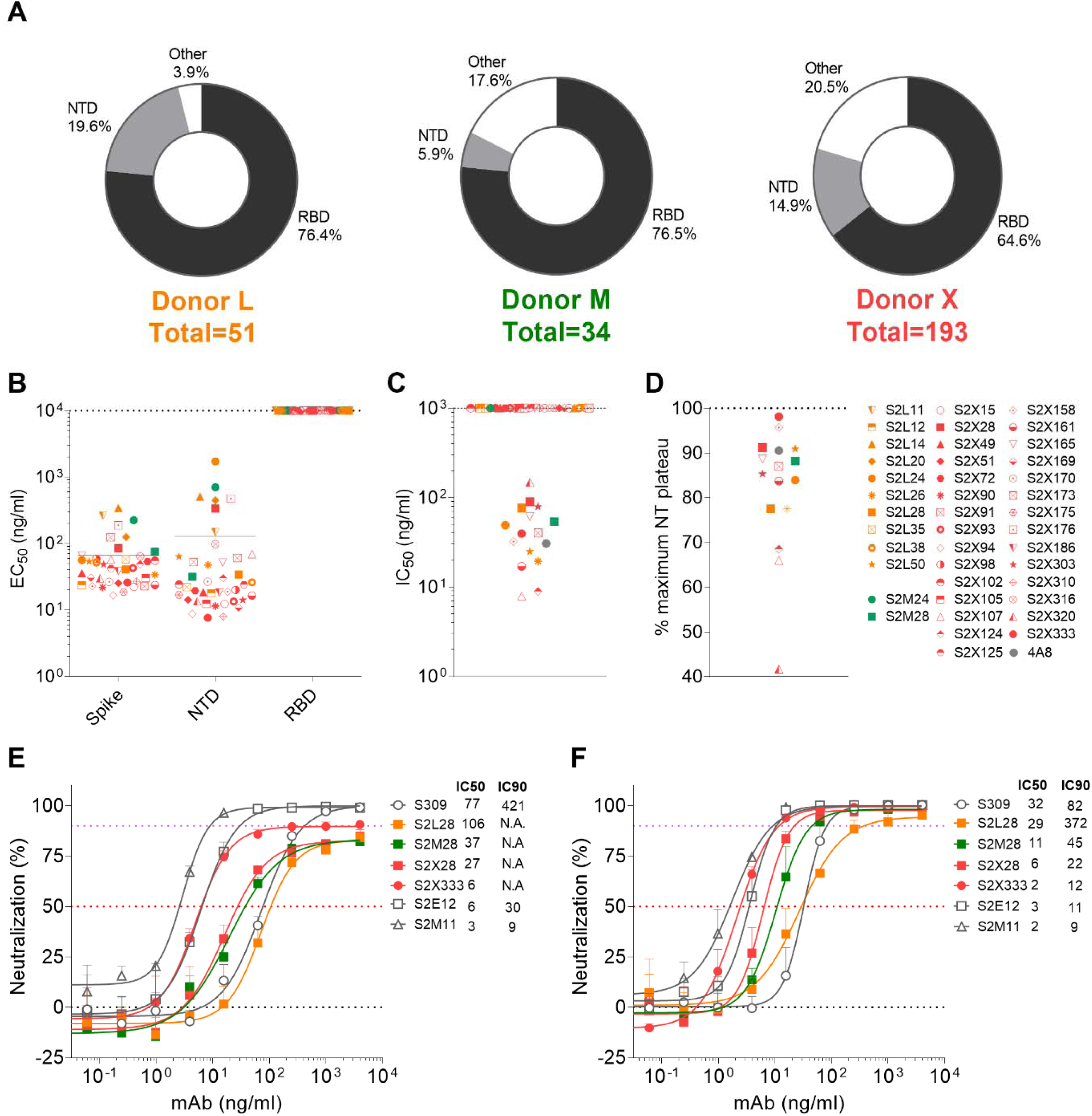
Discovery of potent NTD-specific SARS-CoV-2 neutralizing mAbs from three convalescent individuals. **(A)** Pie charts showing the frequency of mAbs (cloned from IgG^+^ memory B cells) recognizing the SARS-CoV-2 NTD, RBD or other S regions for patients L, M and X. **(B)** Binding of the 41 isolated NTD mAbs to immobilized SARS-CoV-2 S, NTD or RBD analyzed by ELISA. **(C-D)** Neutralization potencies (IC_50_, C) and maximal neutralization plateau (NT, D) of 15 NTD-specific neutralizing mAbs against SARS-CoV-2 S MLV pseudotyped virus. Data are from one out of two independent experiments performed. **(E-F)** Dose-dependent neutralization of selected NTD- and RBD-specific mAbs against authentic SARS-CoV-2-Nluc assessed 6h (at MOI 0.1) (E) or 24h (at M.O.I. of 0.01) (F) after infection. Error bars indicate standard deviation of triplicates.

**Figure S1.**
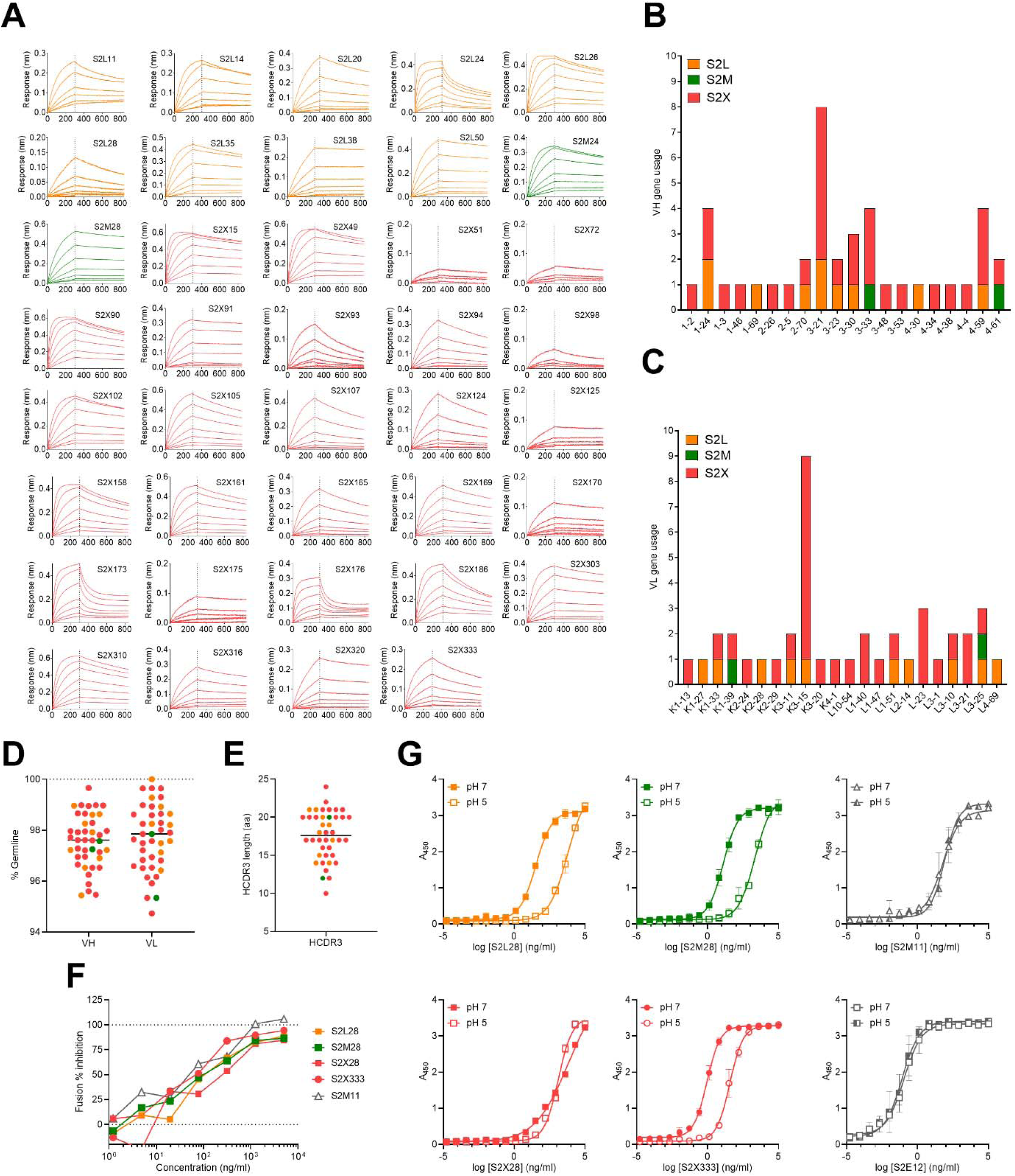
Characteristics of SARS-CoV-2 NTD mAbs. **(A)** Biolayer interferometry binding kinetic analysis of the SARS-CoV-2 NTD to immobilized mAbs. **(B-C)** V gene usage for the heavy (B) and light (C) chains of the NTD mAbs. **(D)** Nucleotide sequence identity of the mAbs isolated relative to the respective V germline genes. **(E)** HCDR3 amino acid length for individual mAbs. **(F)** Cell-to-cell fusion inhibition assay with Vero E6 cells transfected with SARS-CoV-2 S and incubated with varying concentrations of S2L28, S2M28, S2×28, S2×333 or the RBD-specific mAb (S2M11). **(G)** Binding of NTD- and RBD-specific mAbs to immobilized SARS-CoV-2 S at pH7 and pH5 as analyzed by ELISA.

The *in vitro* neutralization activity of the NTD-specific mAbs was subsequently evaluated using a SARS-CoV-2 S pseudotyped murine leukemia virus system (Millet and Whittaker, 2016; Walls et al., 2020c). Out of 41 mAbs, 9 are potent neutralizers (IC_50_ < 50 ng/mL) and 6 are moderate neutralizers (IC_50_ of 50-150 ng/mL) (**Figure 1C and SI Item 3**). The remaining 25 mAbs were non-neutralizing. Most of the mAbs plateaued around 80-90% maximum neutralization in this assay **(Figure 1D and SI Item 2)**. Evaluation of the neutralization potency of a subset of NTD-specific mAbs measured 6 hours post-infection of Vero E6 cells infected with authentic SARS-CoV-2 virus confirmed that these mAbs did not completely block viral entry and instead plateaued at 80-90% neutralization, as opposed to the RBD-specific mAbs S309, S2E12 and S2M11 that achieved 100% neutralization **(Figure 1E)** (Pinto et al., 2020; Tortorici et al., 2020). When the activity was measured at 24 hours post-infection, however, all mAbs tested achieved 95-100% neutralization with a marked enhancement of neutralization potency **(Figure 1F)**. For instance, S2×333 neutralized SARS-CoV-2 with an IC50 of 2 ng/ml and an IC90 of 12 ng/ml, on par with the best-in-class ultrapotent RBD-targeting mAbs S2E12 and S2M11 **(Figure 1F)**. It seems likely that this time course difference may reflect viral spread in the culture.

Previous studies established that SARS-CoV-2 infection of Vero E6 cells proceeds through cathepsin-activated endosomal fusion, as opposed to TMPRSS2-dependent entry which is supposed to occur at the level of the plasma membrane and to be the most relevant route of lung cells infection (Hoffmann et al., 2020a; Hoffmann et al., 2020b; Hoffmann et al., 2020c). Although S2L28, S2M28, S2×28 and S2×333 efficiently block cell-cell fusion **(Figure S1F)**, binding of S2L28, S2M28, and S2×333 to SARS-CoV-2 S was dampened by 2 orders of magnitude at endosomal pH (pH5) compared to neutral pH (pH7). This is expected to reduce their ability to block viral entry at the endosomal level **(Figure S1G)** and might affect differently neutralization after 6 or 24 hours Collectively, these findings indicate that the efficacy of NTD mAbs relies partly on blocking viral entry but also on limiting cell-to-cell spread of the virus.

### NTD-specific neutralizing mAbs delineate an antigenic supersite

To elucidate the mechanism of potent SARS-CoV-2 neutralization by NTD mAbs, we carried out single-particle cryoEM analysis of the SARS-CoV-2 S ectodomain trimer bound to one NTD-specific mAb from each donor – S2L28, S2M28 or S2×333 – in combination with the RBD-specific mAb S2M11. S2M11 was used as it locks the RBDs in the closed state by recognizing a quaternary epitope spanning two adjacent RBDs, thus enabling the use of 3-fold symmetry during reconstruction (Tortorici et al., 2020). 3D classification of the particle images belonging to each dataset revealed the presence of homogeneous ternary complexes with three S2M11 Fabs bound to the RBDs and three bound NTD Fabs radiating from the trimer periphery. We determined reconstructions at 2.6 Å, 2.5 Å and 2.2 Å for the S2L28/S2M11/S, S2M28/S2M11/S and S2×333/S2M11/S complexes **(Figure 2A-I, Figure S2A-C and Table S2)**. We subsequently used local refinement to account for the pronounced conformational dynamics of S2L28, S2M28, and S2×333, and obtained reconstructions at 2.6-3.0 Å resolution for the region comprising the Fab variable domains and their bound epitope on the NTD **(Figure 2A-I, Figure S2A-C and Table S3)**. In parallel, we determined a crystal structure of the SARS-CoV-2 NTD in complex with the S2M28 Fab at 3.0 Å resolution revealing several additional ordered loops and N-linked glycans **(Figure 2 E-H and Table S4)**.

**Figure 2.**
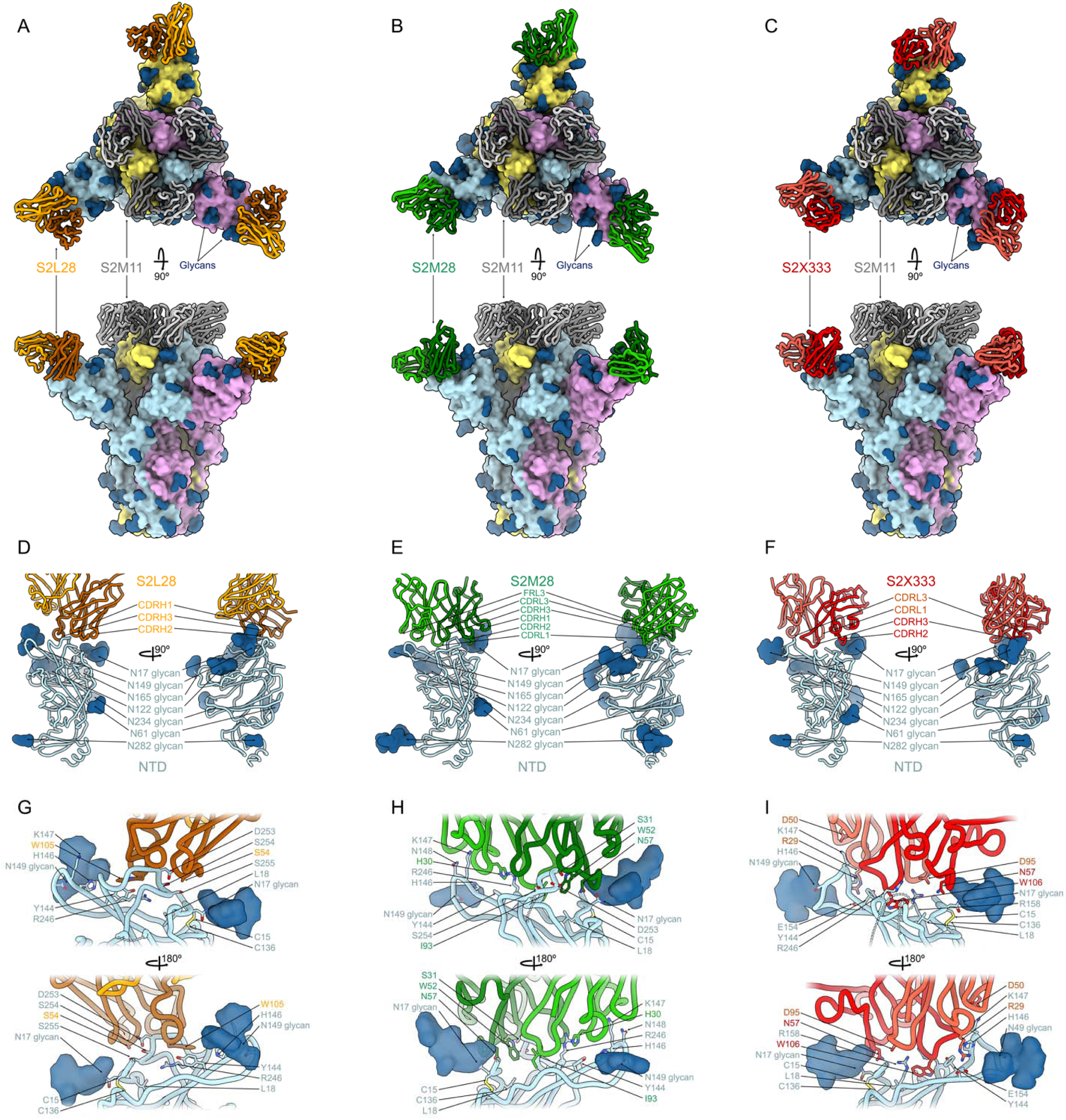
SARS-CoV-2 NTD neutralizing mAbs target the same antigenic supersite. **(A-C)** Ribbon diagrams in two orthogonal orientations of the SARS-CoV-2 S ectodomain trimer (surface) bound to the RBD-specific S2M11 Fab (grey) and to the NTD-targeted Fab S2L28 (A), S2M28 (B) or S2×333 (C). **(D-F)** S2L28 (D), S2M28 (E)and SX333 (F) binding pose relative to the NTD. **(G-I)** Zoomed-in views showing selected interactions of S2L28 (G), S2M28 (H) or S2×333 (I) with the NTD. Missing residues are shown as dotted lines. SARS-CoV-2 S protomers are colored pink, cyan and gold whereas N-linked glycans are rendered as dark blue surfaces.

**Figure S2.**
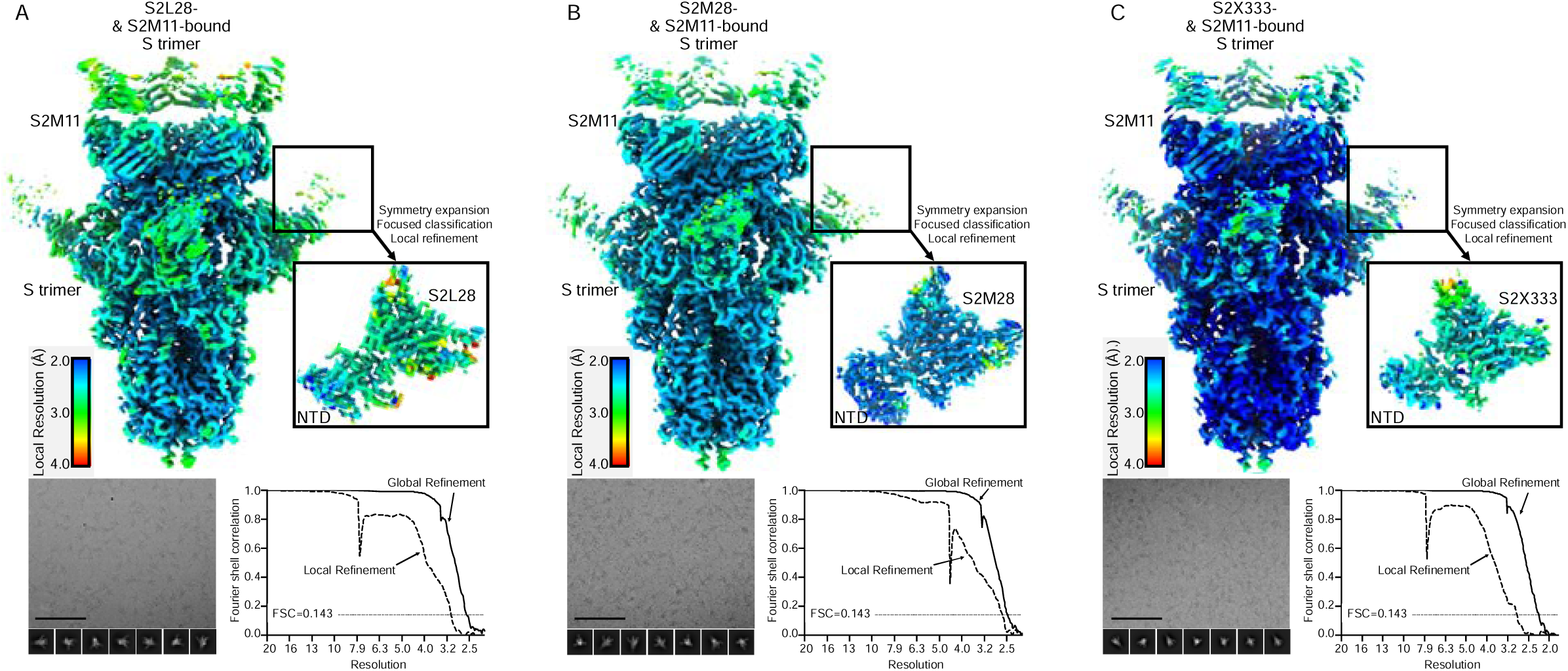
CryoEM data processing of SARS-CoV-2 S bound to S2L28, S2M28, S2×333, S2×28, S2L20, S2×316, or S2M24. Representative electron micrographs and class averages (bottom left of each panel) are shown of SARS-CoV-2 S in complex with the indicated Fabs embedded in vitreous ice (Scale bar: 100 nm). Unsharpened maps colored by local resolution calculated using cryoSPARC (top of each panel) for the S trimer bound to S2M11 Fab and either S2L28 (**A**), S2M28 (**B**), or S2×333 Fab (**C**), as well as the locally refined reconstruction of NTD and NTD-binding Fab.

S2L28, S2M28 and S2×333 contact the N-terminal region (residues 14-20, NTD N-terminus), a β-hairpin formed by residues 140-158 (supersite β-hairpin), and a loop spanning residues 245-264 (supersite loop). These three regions collectively form an antigenic supersite on the pinnacle of the NTD on the side distal to the viral membrane **(Figure 2A-I)**. The epitopes targeted by S2L28, S2M28 and S2×333 are overlapping and flanked by the oligosaccharides at position N17 and N149, which are located at opposite sides of the antigenic supersite **(Figure 2D-I)**. As a stunning example of convergent binding, each mAb uses hydrophobic residues at the tip of the HCDR3 loop to contact the NTD supersite near residue R246. For S2L28, S2M28, and S2×333, this hydrophobic residue is W105, I93, and W106, respectively (**Figure 2G-I**).

Although these Fabs recognize overlapping epitopes within this antigenic supersite, the detailed nature of the interactions formed with the NTD varies. In particular, S2L28 primarily binds to residues in the NTD supersite loop including residue D253 (**Figure 2G**), while S2×333 interacts primarily with the N-terminal region including residue N17 and the supersite β-hairpin including residues E156 and R158 (**Figure 2I**). S2M28 interacts with all the three supersite regions, including residues 146-148 in the supersite β-hairpin, D253 in the supersite loop, and N17 in the N-terminal region (**Figure 2H**). As a result of these distinct interactions, while the S2L28-bound NTD adopts an identical conformation to that observed in the apo S structures (Walls et al., 2020c; Wrapp et al., 2020; Wrobel et al., 2020b), S2M28 and S2×333 induce remodeling of the NTD supersite loop and β-hairpin, respectively (**Figure 2G-I**).

To validate these findings, we analyzed binding of each mAb to purified NTD mutants designed based on the structural data. The R246A substitution decreased binding of S2L28, S2M28 and S2×333 (to various extents), in agreement with the extensive interactions formed by the R246 side chain with aromatic and hydrophobic residues found in the HCDR3 of each mAb **(Figure 3A)**. We found that another neutralizing mAb we isolated (S2×28), as well as the previously described mAb 4A8 (Chi et al., 2020) are also sensitive to the R246A substitution. Removal of the N-linked oligosaccharide at position N17 (T19A) dampened binding to S2M28, S2L28, S2×28, and 4A8 whereas it had minimal effect on S2×333 recognition **(Figure 3A and SI Item 3)**. Abrogation of the N149 oligosaccharide (N149Q), minimally reduced binding for S2×333, S2M28, S2L28, and 4A8, in agreement with the structures showing that this oligosaccharide points away from the epitope/paratope interfaces **(Figure 3A)**. In contrast, S2×28 binding was highly dependent on the N149 glycan, in agreement with our mapping of its epitope to the NTD supersite by cryoEM (**Figure 3A, Figure S2D and SI Item 3**).

**Figure 3.**
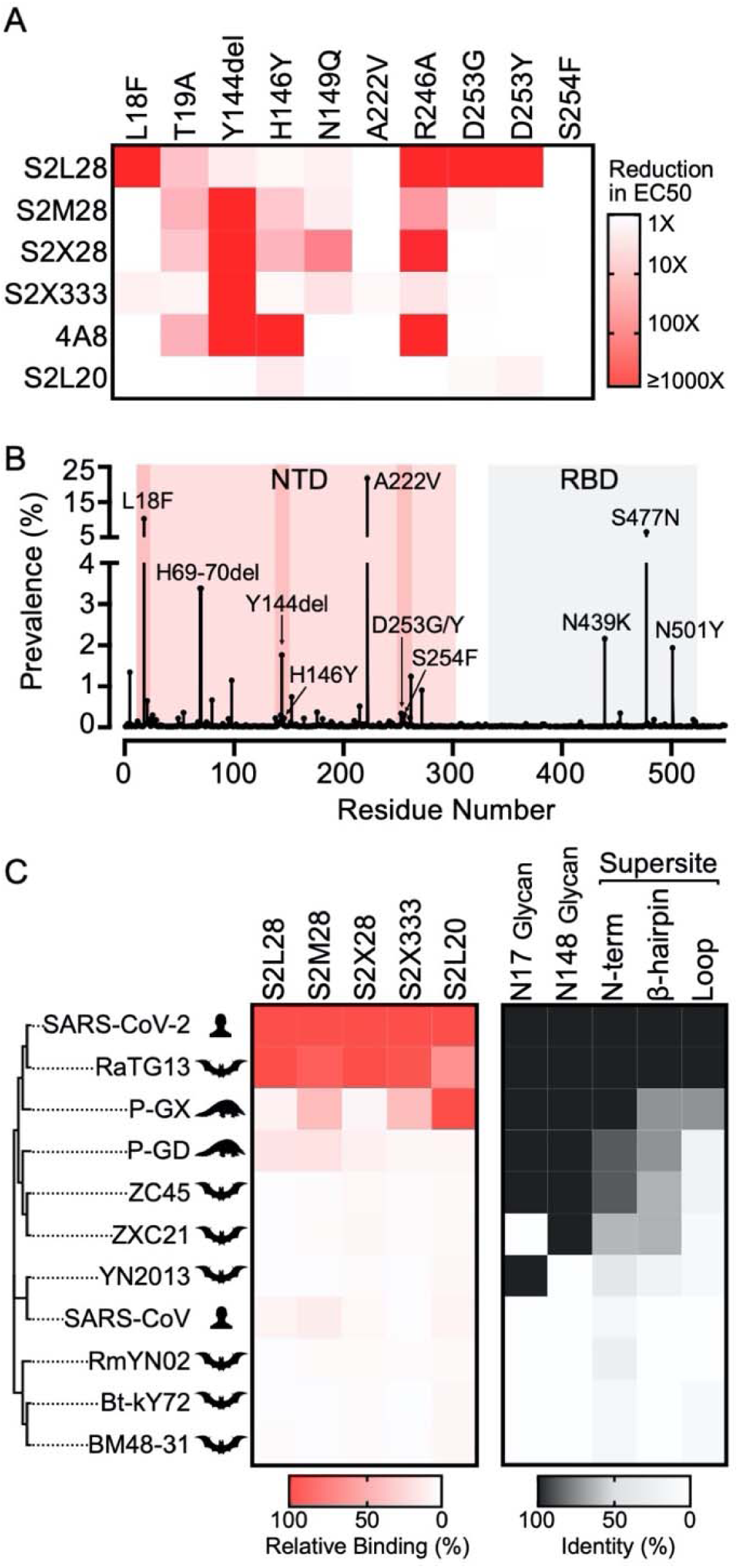
NTD neutralizing mAbs bind to a broad spectrum of SARS-CoV-2 S variants and sarbecovirus S glycoproteins. **(A)** S2L28, S2M28, S2×28, S2×333, 4A8 and S2L20 binding to SARS-CoV-2 NTD variants analyzed by ELISA and displayed as a heat map. Y144del: deletion of residue Y144. **(B)** Prevalence of SARS-CoV-2 S variants among circulating isolates as of December 31^th^ 2020 (290,373 sequences). The NTD and RBD are highlighted with a red and grey backgrounds, respectively; the NTD supersite residues are highlighted by darker red background shading. **(C)** Heatmap of NTD mAb binding to a panel of sarbecovirus S glycoproteins expressed at the surface of ExpiCHO cells analyzed by flow-cytometry. Binding is expressed as mean fluorescence intensities relative to SARS-CoV-2 S binding for each mAb. A coronavirus NTD cladogram is shown on the left. The conservation of relevant glycans and NTD supersite regions is also shown (right).

**Figure S3.**
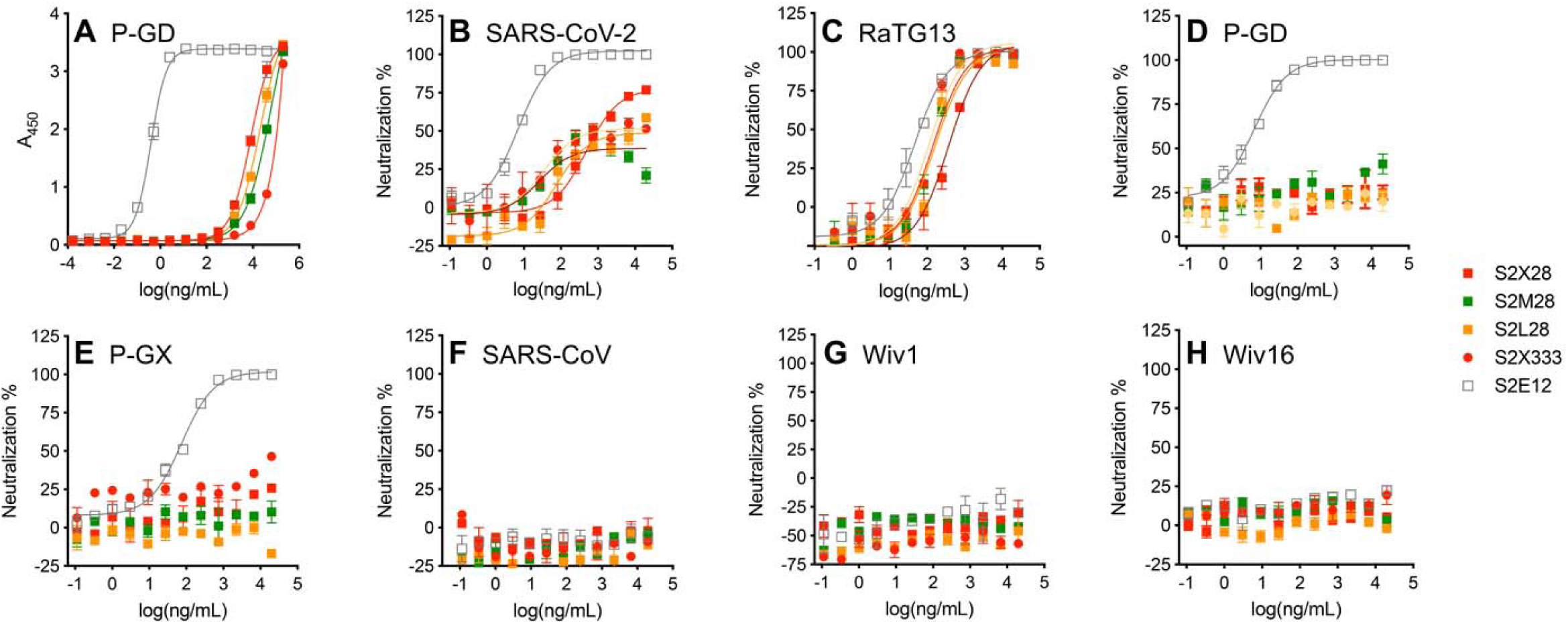
NTD neutralizing mAbs inhibit S-mediated entry of the closely related RaTG13 but not of more distant sarbecoviruses. **(A)** Binding of NTD- and RBD-specific mAbs to immobilized P-GD S ectodomain trimer analyzed by ELISA. **(B-H)** VSV pseudovirus neutralization assays in the presence of varying concentrations of the NTD-specific mAbs S2L28, S2M28, S2×58, S2×333 or the RBD-specific mAb S2E12.

CryoEM maps of S2L28-, S2M28-, S2×28-, and S2×333-bound SARS-CoV-2 S delineate an antigenic supersite recognized by potent NTD-specific neutralizing mAbs isolated from COVID-19 individuals. Likewise, the previously described mAb 4A8 (Chi et al., 2020) as well as the recently described FC05 (Zhang et al., 2020) and CM25 (Voss et al., 2020), bind to the same NTD antigenic supersite to neutralize SARS-CoV-2. Collectively, these results show that NTD-specific neutralizing mAbs have adopted a convergent mechanism of viral inhibition through the use of multiple V genes **(Table S1)**, which points to a key role of this region of the NTD in viral neutralization and putatively also infection.

### NTD mAbs bind to a broad spectrum of SARS-CoV-2 isolates and neutralize RaTG13

Analysis of the 290,373 SARS-CoV-2 genome sequences deposited in GISAID as of December 31^st^ 2020 reveals a larger number of prevalent mutations and deletions in the NTD than other regions of the S glycoprotein **(Figure 3B)**. Besides the A222V variant, which has a prevalence exceeding 21% (Hodcroft et al., 2020), we identified several variants within the NTD antigenic supersite **(Figure 3A-B)**, which based on the structural data, might impact mAb binding. These variants are L18F, a Y144 deletion, H146Y, D253G/Y and S254F with prevalence of 10% (39 countries), 1.7% (37 countries), 0.2% (15 countries), 0.3/0.005% (9 countries) and 0.1% (27 countries), respectively.

Assessment of mAb binding to purified SARS-CoV-2 NTD harboring these mutations revealed that all neutralizing NTD mAbs bind efficiently to the A222V NTD variant, indicating these mAbs would not be affected by this common SARS-CoV-2 mutation. L18F or D253G/Y variants abrogated binding only to S2L28, which is relevant considering L18F is the third most prevalent mutant sequenced to date **(Figure 3B)**. Conversely, the Y144 deletion abrogated binding to S2M28, S2×28, S2×333, and 4A8, but not S2L28 **(Figure 3A and SI Item 3)**. In addition, the H146Y mutant reduced binding to S2M28, S2×28, and especially 4A8 **(Figure 3A and SI Item 3)**. Binding of the non-neutralizing S2L20 NTD-specific mAb was not affected by any of these variants, confirming retention of proper folding of the purified NTD mutants **(Figure 3A and SI Item 3)**. Collectively these data show that S2M28, S2×28, S2×333, and 4A8 have a similar reactivity profile to these NTD mutants, whereas S2L28 has a distinct and complementary pattern. These data and those used to validate the structural analysis suggest that several currently circulating variants may escape from neutralizing mAbs targeting the antigenic supersite. Indeed, the L18F and R246I mutations have been identified in the 501Y.V2 lineage, whereas a Y144 deletion is found in the B.1.1.7 lineage that have recently exhibited a marked rise in frequency in South Africa and the UK, respectively (Davies et al., 2020; Tegally et al., 2020).

To investigate the cross-reactivity of NTD-specific mAbs with sarbecovirus S glycoproteins, we probed individual mAb binding to ExpiCHO cells expressing full-length S from representative members of the three clades of the sarbecovirus subgenus. All mAbs tested bound to RaTG13 S, which is the most closely related S glycoprotein and NTD to SARS-CoV-2 S **(Figure 3C and SI Item 4)**. S2×333 and S2M28 weakly bound to Pangolin Guanxi 2017 (P-GX) S, whereas S2L28 and S2M28 weakly bound to Pangolin Guangdong 2019 (P-GD) S **(Figure 3C and SI Item 4)**. Consistently, weak mAb binding to P-GD S was confirmed by ELISA for S2L28, S2M28, S2×333 and S2M24 (**Figure S3A**). We next evaluated the neutralization breadth of S2L28, S2M28, S2×28 and S2×333 against SARS-CoV-2- and SARS-CoV-related sarbecoviruses using a vesicular stomatitis virus (VSV) pseudotyped neutralization assay. All mAbs tested neutralized RaTG13 S pseudotypes with comparable potency to SARS-CoV-2 S pseudotyped virus but none of them neutralized the more distantly related P-GX S, P-GD S, SARS-CoV S, WIV1 S or WIV16 S pseudotyped viruses **(Figure S3B-H)**. These findings are in line with the low conservation of the NTD supersite in phylogenetically distant viruses and indicate that NTD mAbs could be useful countermeasures against viruses closely related to SARS-CoV-2, but not against divergent viruses. Both species-specific functions and host immunity may play a role in the antigenic diversification of this region compared to other S domains that are more conserved.

### SARS-CoV-2 S neutralization escape mutants reveal unconventional mechanisms of NTD mAb resistance

To investigate potential viral escape from neutralization mediated by the potent neutralizing NTD mAbs, we passaged a replication competent VSV-SARS-CoV-2 chimera in which the native glycoprotein was replaced by the SARS-CoV-2 S Wuhan-1 gene in the presence of S2L28, S2M28, S2×28 and S2×333, as previously described (Case et al., 2020; Liu et al., 2020b). Selective pressure provided by each mAb led to emergence of several escape mutants that clustered within the antigenic supersite i.e. the NTD N-terminus, the supersite β-hairpin, and the supersite loop as well as surrounding residues **(Figure 4A-B and SI Item 3 and 5)**.

**Figure 4.**
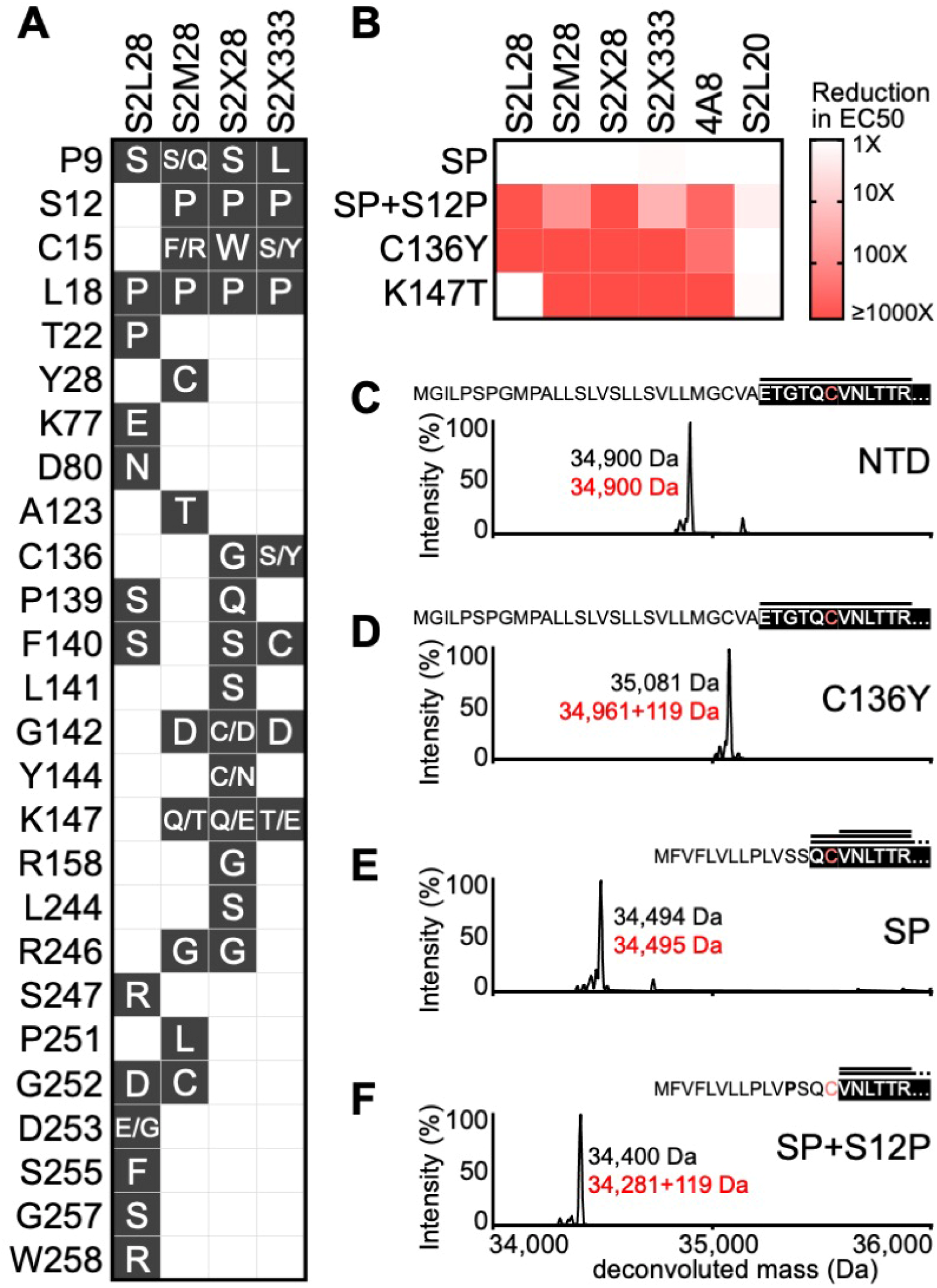
Analysis of SARS-CoV-2 S neutralization escape mutants. **(A)** Escape mutations selected with each mAb (black background white font) and residues that remained unchanged (white background). (**B**) S2L28, S2M28, S2×28, S2×333, 4A8 and S2L20 binding to selected SARS-CoV-2 NTD escape mutants analyzed by ELISA and displayed as a heat map. The native signal peptide was introduced (SP) and also introduced along with the S12P mutation (SP+S12P). **(C-F)** Deconvoluted mass spectra of purified NTD constructs, including the NTD construct with an optimized signal peptide (**C**, NTD), the NTD construct with an optimized signal peptide and the C136Y mutation (**D**, NTD), the NTD construct with the native signal peptide (**E**, NTD), and the NTD construct with the native signal peptide and the S12P mutation (**F**, NTD). The empirical mass (black) and theoretical mass (red) are shown beside the corresponding peak. Additional 119 Da were observed for the C136Y and SP+S12P NTDs corresponding to cysteinylation of the free cysteine residue in these constructs (as L-cysteine was present in the expression media). The cleaved signal peptide (black text white background) and subsequent N-terminal sequence (white text black background) are also shown; C15 is highlighted in light red. Peptides identified by MS/MS analysis consistent with the mass of N-terminal peptides are shown above the N-terminal sequence (black horizontal lines).

**Figure S4.**
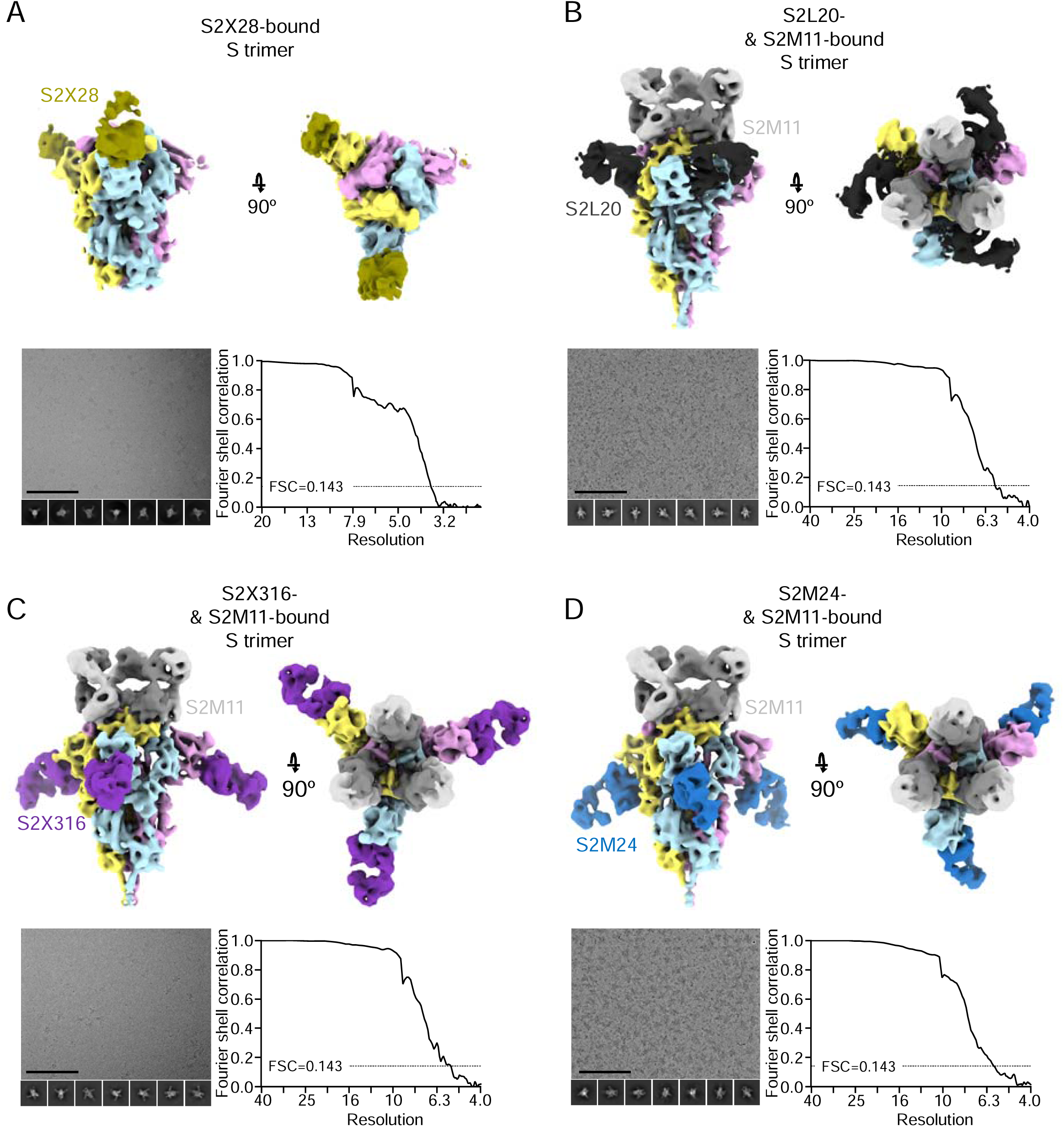
CryoEM data processing of SARS-CoV-2 S bound to S2×28, S2L20, S2×316, or S2M24. Representative electron micrographs and class averages (bottom left of each panel) are shown of SARS-CoV-2 S in complex with the indicated Fabs embedded in vitreous ice (Scale bar: 100 nm). (**A**) 6 Å low-pass filtered map of the SARS-CoV-2 2P DS S trimer (McCallum et al., 2020) bound to S2×28 (gold). (**B-D**) Sharpened maps of the S trimer bound to S2M11 Fabs and S2L20 (**B**, grey), S2×316 (**C**, purple), or S2M24 Fabs (**D**, sky blue). The corresponding Fourier shell correlation curves (bottom right of each panel) are shown with the 0.143 cutoff indicated by horizontal dashed lines.

All four mAb selections share many escape mutations but with a distinct pattern for S2L28, in agreement with the structural observations indicating distinct residues are important for S2L28 recognition. Similarly to what was described previously for RBD mAbs (Starr et al., 2020a; Starr et al., 2020b), all viral escape mutants resulted from a single non-synonymous nucleotide mutation. For instance, S2L28 selected for the D253E and D253G substitutions whereas S2×28 selected for the Y144C and Y1444N substitutions. Furthermore, both S2M28 and S2×28 selected for the R246G substitutions. Deletion of residue Y144 or the R246I/T/K and D253G/Y mutations have been detected in SARS-CoV-2 clinical isolates demonstrating the ability of this system to recapitulate variants at naturally occurring S residues.

Besides residue substitutions that are expected to impact the epitope/paratope interface directly, we observed escape mutants in the signal peptide region and at cysteine residues 15 and 136. The latter two residues are jointly engaged in a disulfide bond that staples the NTD N-terminus against the galectin-like β-sandwich **(Figure 2G-I)**. Alteration of this bond (via C136Y substitution) abrogates recognition by S2L28, S2M28, S2×28, S2×333 and 4A8 **(Figure 4B and SI Item 3)**, likely via dislodging this key supersite component, explaining the observed phenotype. These data further suggest that this mechanism of escape is likely to be effective with all mAbs targeting the NTD antigenic supersite.

The predicted SARS-CoV-2 S signal peptide cleavage site occurs between residues T13 and Q14. Signal peptide cleavage site prediction using SignalP5.0 (Almagro Armenteros et al., 2019) suggests that the S12P escape mutation alters cleavage to occur between residues C15 and V16, thereby eliminating the C15-C136 disulfide bond – similarly to escape mutants at positions C15 and C136. To test the effect of S12P with purified NTD, we introduced the native signal peptide with and without S12P. Similar to direct disruption of the C15-C136 disulfide bond, the S12P signal peptide mutation was found to markedly reduce binding to S2L28, S2M28, S2×28, S2×333 and 4A8 **(Figure 4B and SI Item 3)**. Mass spectrometry analysis of the S12P mutant was consistent with signal peptide cleavage occurring immediately after residue C15 **(Figure 4C-F)**. These data warrant closer monitoring of signal peptide variants and their involvement in immune evasion since the S12F mutation has been found in SARS-CoV-2 clinical isolates (0.1% prevalence) and is predicted to have a similar effect to the S12P substitution characterized here.

### Definition of a SARS-CoV-2 NTD antigenic map

To understand the markedly different neutralization potencies across the 41 NTD-specific mAbs identified, we carried out competition biolayer interferometry binding assays using recombinant SARS-CoV-2 S. The data indicated that the mAbs recognize six distinct antigenic sites, which we designated i, ii, iii, iv, v and vi. Most mAbs clustered within antigenic sites i and iii whereas sites ii, iv, v and vi each accounted for only one or a small number of mAbs from our panel **(Figure 5A and SI Item 6)**. All potently neutralizing mAbs competed for binding to the NTD site i, indicating they recognize overlapping epitopes within the structurally identified antigenic supersite **(Figure 1B-E, Figure 2 and Figure 5A)**.

**Figure 5.**
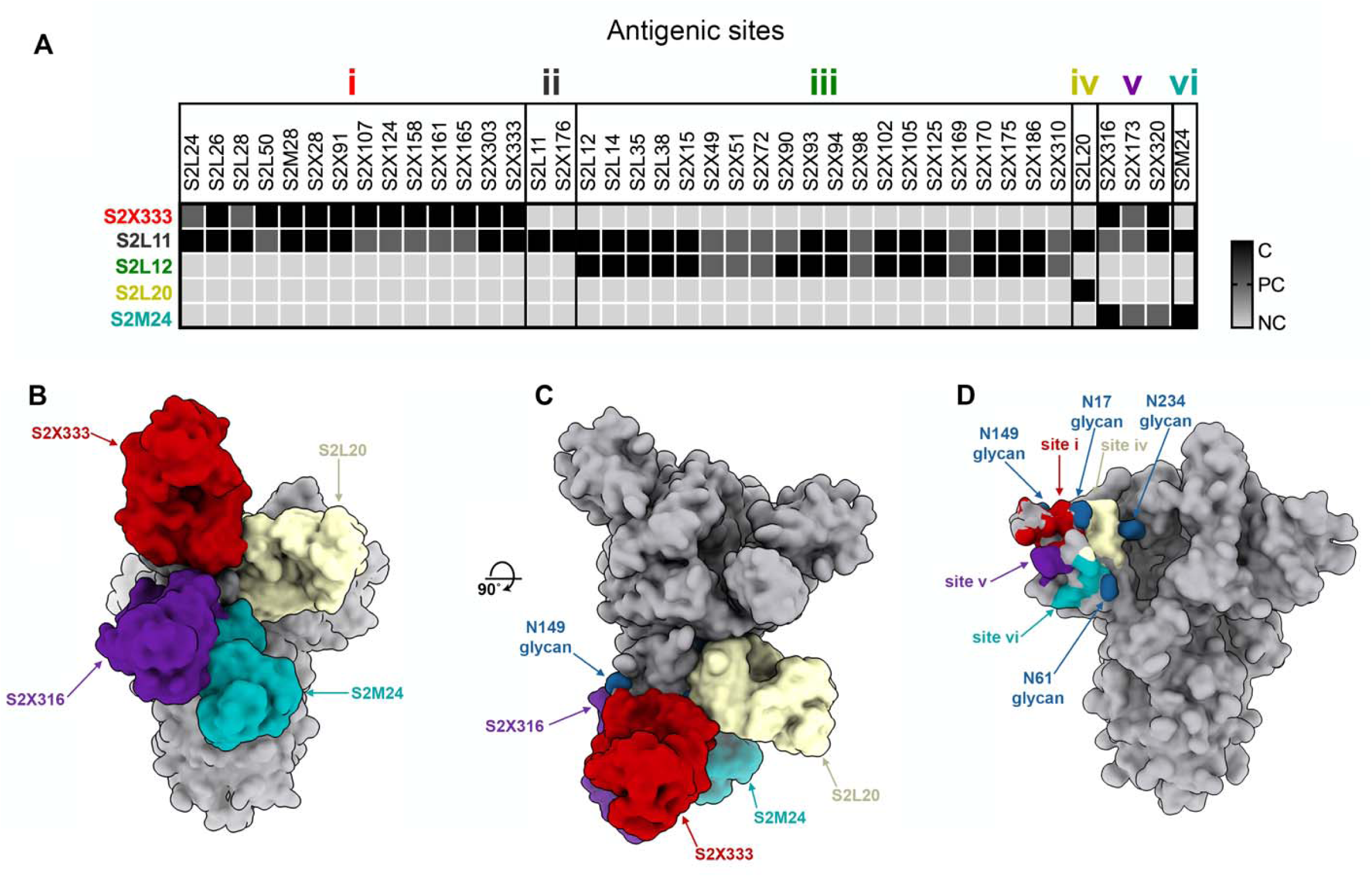
The SARS-CoV-2 NTD comprises multiple antigenic sites. **(A)** Epitope binning of the 41 NTD-specific mAbs isolated led to the identification of 6 antigenic sites based on competition binding assays using biolayer interferometry. NC: no competition; PC: partial competition; C: competition **(B-C)** Composite model of the SARS-CoV-2 S trimer viewed along two orthogonal orientations with four bound Fab fragments representative of antigenic site i (S2×333), site iv (S2L20), site v (S2×316) and site vi (S2M24). **(D)** Footprints of the antigenic sites identified structurally are shown along with neighboring N-linked glycans (blue spheres).

**Figure S5.**
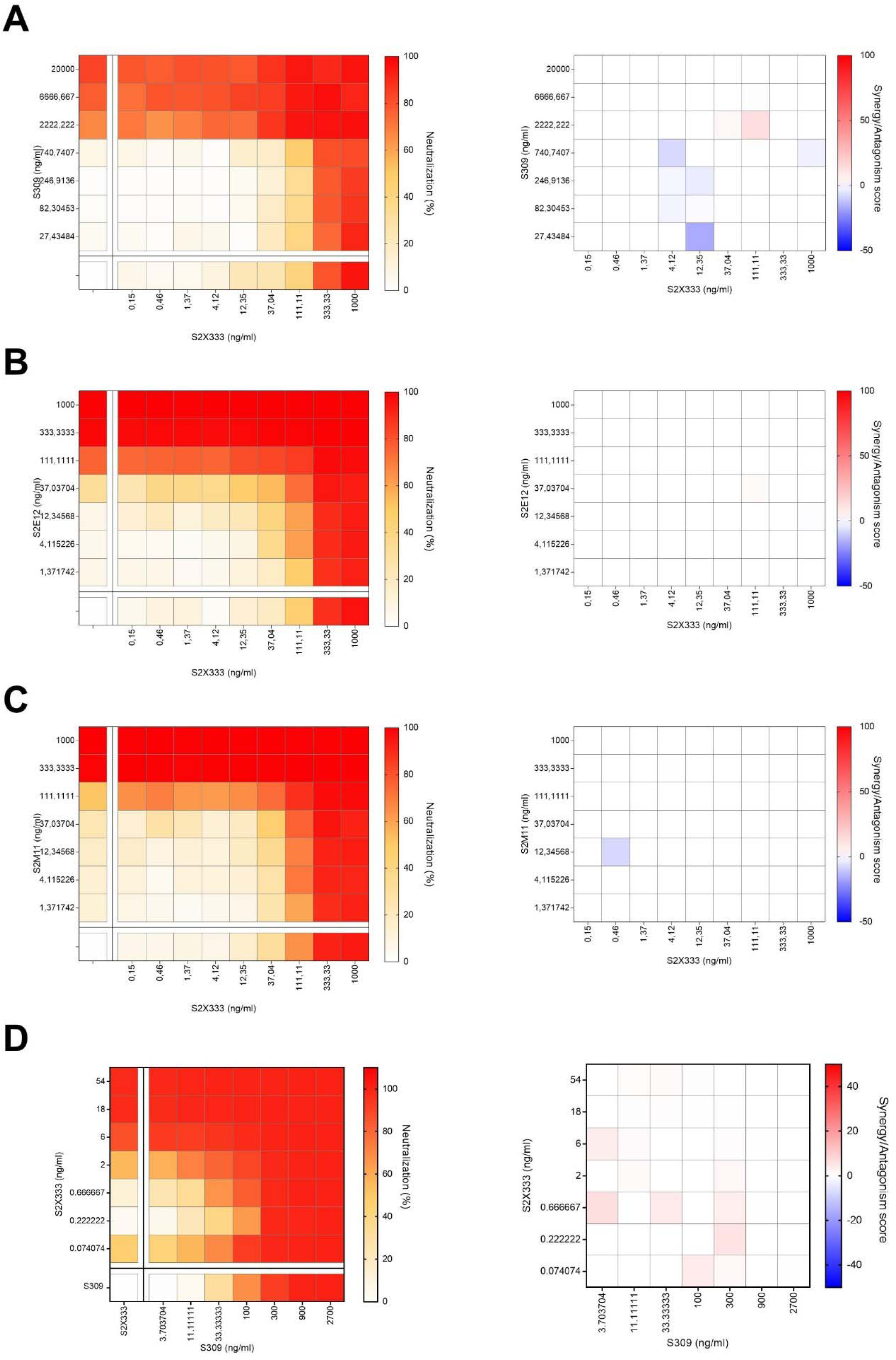
Neutralizing activity of SARS-CoV-2 NTD- and RBD-targeting antibody cocktail. **(A-C)** SARS-CoV-2-MLV pseudotypes neutralization (left) and synergy score (right) measured combining S2×333 with the RBD-targeting mAb S309 (A), S2E12 (B), or S2M11 (C). **(D)** Neutralization matrix to assess the synergistic activity of S2×333 and S309 mAb cocktails *in vitro* with authentic SARS-CoV-2-Nluc. Data for authentic SARS-CoV-2-Nluc are from one representative experiment performed in triplicate each.

To delineate the different NTD antigenic sites, we mapped the epitopes targeted by representative mAbs through determination of cryoEM structures of S2M24, S2×316 or S2L20 in complex with SARS-CoV-2 S, together with S2M11 to expedite structural determination at 6-8Å resolution **(Figure S4 and Table S2)**. S2L20 (site iv) binds to an epitope flanked by glycans at positions N17, N61 and N234 located within the NTD side which faces the RBD belonging to the same protomer **(Figure 5B-D)**. S2M24 (site vi) recognizes the viral membrane proximal side of the NTD which is devoid of glycans except for the presence of the N61 oligosaccharide on the edge of the epitope **(Figure 5B-D)**. S2M24 recognizes a similar antigenic site as the one described for the polyclonal Fabs COV57 isolated directly from COVID-19 patient sera (Barnes et al., 2020b). S2×316 (site v) interacts with an epitope residing nearby glycans N74 and N149 at the peripheral-most part of the NTD, located in between antigenic sites i and vi **(Figure 5B-D)**. As all NTD-specific mAbs with neutralizing activity target the structurally identified antigenic supersite site (site i), our data points to the existence of a site of vulnerability that is harnessed by the immune system of multiple individuals.

### Mechanism of action of NTD-specific neutralizing mAbs

To understand NTD-specific mAb-mediated inhibition of viral entry, we evaluated the ability of these mAbs to block ACE2 binding as this step correlates with neutralization titers in SARS-CoV-2 exposed individuals (Piccoli et al., 2020). None of the site i-targeting mAbs (S2L28, S2M28, S2×28, and S2×333) blocked binding of SARS-CoV-2 S to immobilized human recombinant ACE2 as measured by biolayer interferometry **(Figure 6A)**, excluding interference with engagement of the main entry receptor as mechanism of action. Moreover, these mAbs did not promote shedding of the S_1_ subunit from cell-surface-expressed full-length SARS-CoV-2 S **(Figure 6B)**, ruling out the possibility of premature S triggering, as previously shown for a SARS-CoV and several SARS-CoV-2 RBD-specific mAbs (Huo et al., 2020; Piccoli et al., 2020; Walls et al., 2019; Wec et al., 2020; Wrobel et al., 2020a). However, S2L28, S2M28, S2×28, and S2×333 blocked SARS-CoV-2 S-mediated cell-cell fusion, indicating they could either prevent interaction with an auxiliary receptor, proteolytic activation or membrane fusion **(Figure S1G)**.

**Figure 6.**
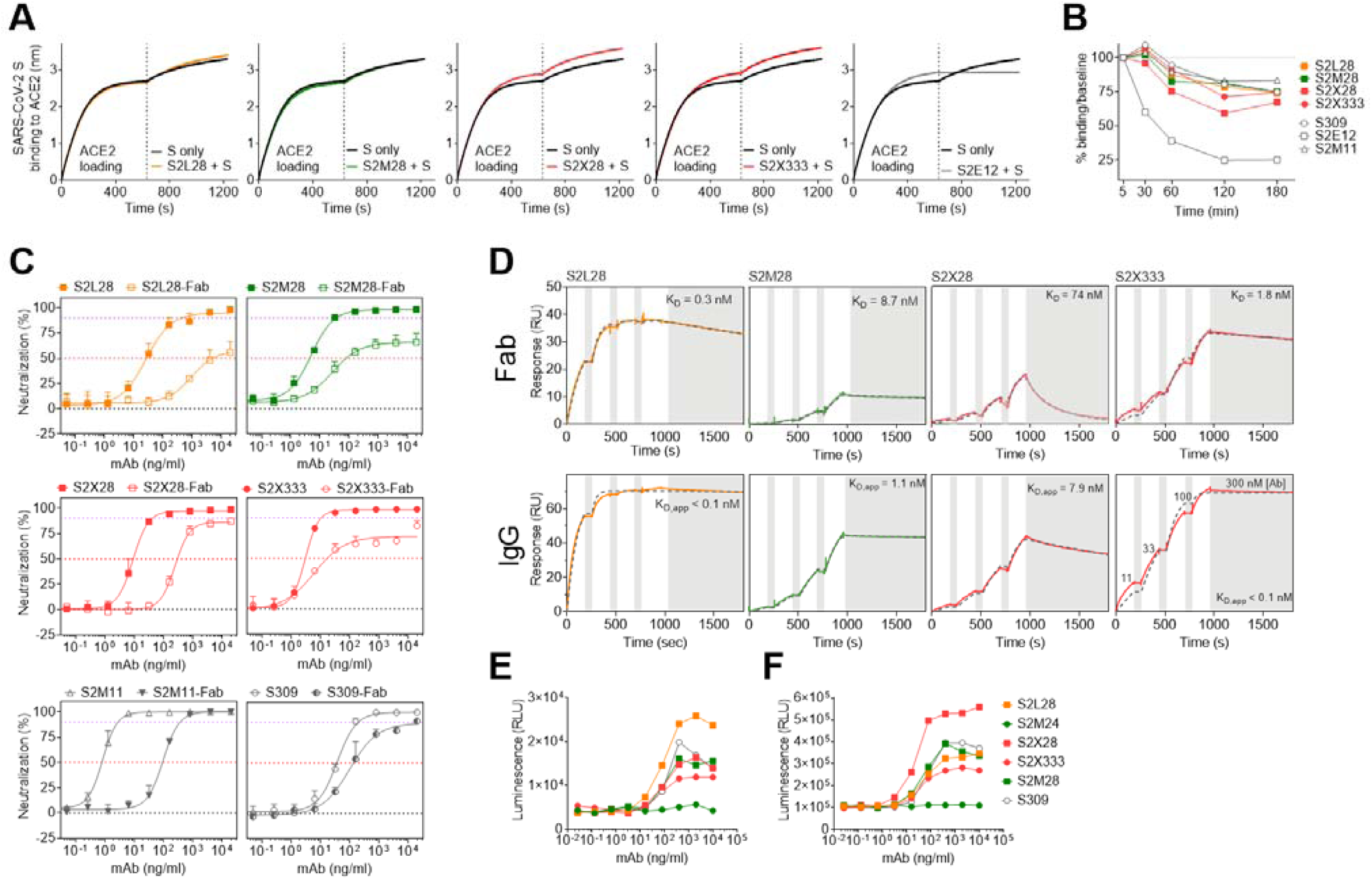
Mechanism of action of NTD-specific neutralizing mAbs. **(A)** Competition of S2L28, S2M28, S2×28, and S2×333 with ACE2 to bind to SARS-CoV-2 S as measured by biolayer interferometry. ACE2 was immobilized at the surface of the biosensors before incubation with S alone or in complex with the mAbs. The vertical dashed line indicates the start of the association of S/mAb complex or free S with solid-phase ACE2. The anti-RBD S2E12 mAb was included as positive control. (**B)** mAb-mediated S_1_ subunit shedding from cell-surface expressed SARS-CoV-2 S as determined by flow-cytometry. The anti-RBD S2M11 and S2E12 mAbs were included as negative and positive controls, respectively. (**C)** Neutralization of authentic SARS-CoV-2 (SARS-CoV-2-Nluc) by S2L28, S2M28, S2×28, and S2×333 IgG or Fab. S309 and S2M11 Fab and IgG were also included as controls. Symbols are means ± SD of triplicates. Dotted lines indicate IC_50_ and IC_90_ values. **(D)** SPR analysis of mAbs binding to the SARS-CoV-2 S ectodomain trimer. Gray dashed line indicates a fit to a 1:1 binding model. The equilibrium dissociation constants (K_D_) or apparent equilibrium dissociation constants (K_D_, app) are indicated. White and gray stripes indicate association and dissociation phases, respectively. **(E-F)** Activation of FcγRIIa H131 (E) and FcγRIIIa V158 (F) induced by NTD-specific mAbs. The anti-RBD mAb S309 was included as positive control.

To assess whether steric hindrance plays a role in the neutralization activity of S2L28, S2M28, S2×28, and S2×333, we evaluated the neutralization potency of each mAb, in both Fab and IgG formats, against authentic SARS-CoV-2-Nluc **(Figure 6C)**. NTD-specific Fabs displayed a marked potency reduction, both in terms of IC_50_ values and maximal neutralization plateau reached **(Table S1)**, as compared to IgGs, possibily due to reduced avidity as observed by surface plasmon resonance (**Figure 6D**). Since the Fabs could still partially neutralize SARS-CoV-2, at least part of the observed neutralization activity results from direct interaction with their respective epitopes. It is possible that NTD-specific mAb-mediated neutralization further relies on the steric hindrance provided by Fc positioning, similar to what was observed for anti-hemagglutinin influenza A virus neutralizing mAbs (Xiong et al., 2015).

We next examined potential additive, antagonistic or synergistic effects of NTD- and RBD-targeting mAbs, as mAb synergy was previously described for SARS-CoV and SARS-CoV-2 neutralization (Pinto et al., 2020; ter Meulen et al., 2006). Cocktails of S2×333 with S309, S2E12, or S2M11 additively prevented entry of SARS-CoV-2 S-MLV pseudotyped virus in Vero E6 cells **(Figure S5A-C)**. This additive effect was also observed between S2×333 and S309 using authentic SARS-CoV-2 at 24 hours post-infection in Vero E6 cells **(Figure S5D)**. These results are consistent with RBD- and NTD-targeting mAbs mediating inhibition by distinct mechanisms and demonstrate that they could be used as cocktails for prophylaxis or therapy.

Since Fc-mediated effector functions contribute to protection by promoting viral clearance and anti-viral immune responses *in vivo* (Bournazos et al., 2020; Bournazos et al., 2016; Schäfer et al., 2021; Winkler et al., 2020), we evaluated the ability of site i-targeting mAbs to trigger activation of FcγRIIa and FcγRIIIa as a proxy for Ab-dependent cellular phagocytosis (ADCP) and Ab-dependent cellular cytotoxicity (ADCC), respectively. S2L28, S2M28, S2×28, and S2×333 promoted dose-dependent FcγRIIa and FcγRIIIa-mediated signaling to levels comparable to those of the highly effective mAb S309 (Pinto et al., 2020) **(Figure 6E-F)**. In contrast, the non-neutralizing site vi-targeting S2M24 mAb did not promote FcγR-mediated signaling, possibly due to the different orientation relative to the membrane of the effector cells in comparison to site i-specific mAbs **(Figure 6E-F)**. These findings suggest that besides their neutralizing activity, mAbs recognizing site i can exert a protective activity via promoting Fc-mediated effector functions.

### NTD neutralizing mAbs protect against SARS-CoV-2 challenge in hamsters

To evaluate the protective efficacy of NTD-directed mAbs against SARS-CoV-2 challenge *in vivo*, we selected the S2×333 mAb for a prophylactic study in a Syrian hamster model (Boudewijns et al., 2020). The mAb was administered at 4 and 1 mg/kg via intraperitoneal injection 48 hours before intranasal SARS-CoV-2 challenge. Four days later, lungs were collected for the quantification of viral RNA and infectious virus titers. Prophylactic administration of S2×333 decreased the amount of viral RNA detected in the lungs by ∼3 orders of magnitude, compared to hamsters receiving a control mAb **(Figure 7A)** and completely abrogated viral replication in the lungs of most animals at both doses tested **(Figure 7B)**. Although all animals had similar serum mAb concentrations within each group, no reduction in the amount of viral RNA or infectious virus was observed for one hamster at each dose compared to those administered with a control mAb **(Fig. 7C-D)**. Based on the aforementioned variability and mutation tolerance of the SARS-CoV-2 NTD, we hypothesize that S2×333 escape mutants might have been selected in these animals. Overall, these data show that low doses of anti-NTD mAbs provide remarkable prophylactic activity *in vivo*, comparable to best-in-class RBD-specific mAbs S2E12 and S2M11 (Tortorici et al., 2020), consistent with their potent *in vitro* neutralizing activity. We anticipate that the protection efficacy of S2×333 (and related NTD mAbs) would be further enhanced in humans due to an optimal matching with human Fcγ receptors.

**Figure 7.**
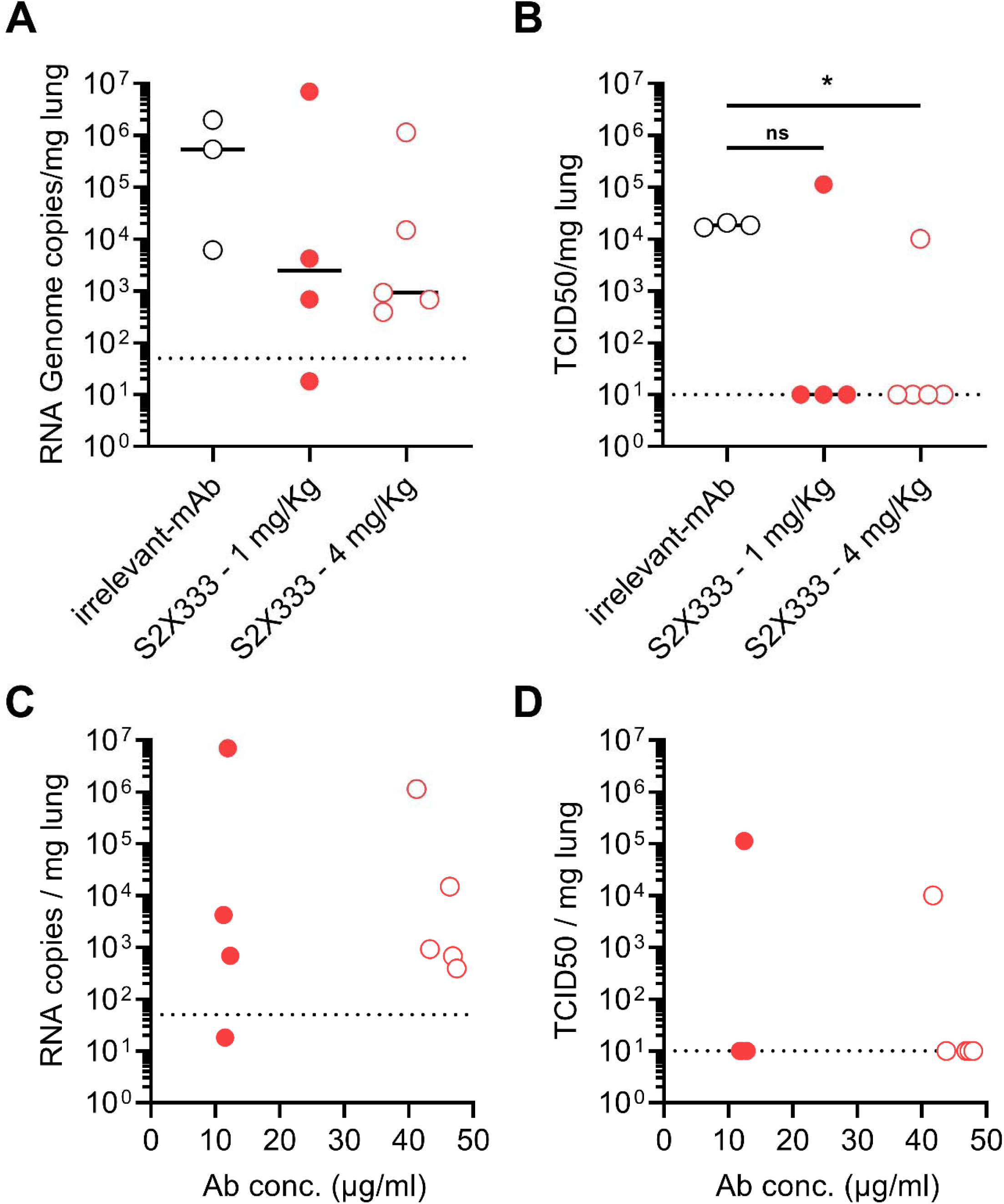
S2×333 provides robust in vivo protection against SARS-CoV-2 challenge. Syrian hamsters were injected with the indicated amount of mAb 48 h before intra-nasal challenge with SARS-CoV-2. **(A)** Quantification of viral RNA in the lungs 4 days post-infection. **(B)** Quantification of replicating virus in lung homogenates harvested 4 days post infection using a TCID_50_ assay. **(C-D)** Viral RNA loads and replicating virus titers in the lung 4 days post infection plotted as a function of serum mAb concentrations before infection (day 0).

## Discussion

The data herein demonstrate that neutralizing NTD-targeting mAbs represent a key aspect of immunity to SARS-CoV-2 and account for 5-20% of SARS-CoV-2 S-specific mAbs cloned from memory B cells isolated from the PBMCs of three COVID-19 individuals. Analysis of a large panel of neutralizing and non-neutralizing mAbs led us to define an antigenic map of the heavily glycosylated SARS-CoV-2 NTD, in which 6 antigenic sites (i-vi) were identified. Importantly, we found that all the neutralizing mAbs from the three donors investigated targeted the same antigenic supersite (site i). The neutralizing mAbs described here along with the mAbs 4A8 (Chi et al., 2020), FC05 (Zhang et al., 2020) and CM25 (Voss et al., 2020), which also target this antigenic supersite, use various germline V genes to recognize overlapping epitopes, thereby providing examples of convergent solutions to NTD-targeted mAb neutralization. We show that a highly potent NTD mAb provides prophylactic protection against SARS-CoV-2 challenge of Syrian hamsters showcasing that this class of mAbs can be a critical barrier to infection and are promising clinical candidates against SARS-CoV-2. Likewise, previous work on MERS-CoV established that the NTD is the target of several potent and protective neutralizing mAbs and an attractive target for vaccine design (Chen et al., 2017; Jiaming et al., 2017; Wang et al., 2018; Wang et al., 2019; Zhou et al., 2019).

The emergence and fixation of SARS-CoV-2 D614G yielded a virus mutant with enhanced ACE2 binding, transmissibility and replication but without marked changes of antigenicity (Hou et al., 2020; Korber et al., 2020; Plante et al., 2020; Yurkovetskiy et al., 2020). In contrast, we observed that several variants detected in circulating SARS-CoV-2 clinical isolates reduced or abrogated recognition by some NTD-specific neutralizing mAbs. For instance, the L18F substitution has a prevalence of 10% among sequenced SARS-CoV-2 genomes, is present in the 501Y.V2 lineage initially identified in South Africa (Tegally et al., 2020), and would escape S2L28-mediated neutralization. A deletion of residue Y144 is found in 1.7% of circulating SARS-CoV-2 clinical isolates, including the B.1.1.7 lineage that was originally detected in the UK before being identified in numerous other countries (Davies et al., 2020). This variant circumvents recognition by S2M28, S2×28, S2×333 and 4A8. We identified additional key residues for binding of neutralizing mAbs, such as S12 and R246, using viral escape selection and structural analysis for which natural variants are reported and are thus expected to participate in neutralization escape. The finding that multiple circulating SARS-CoV-2 variants map to the NTD, including several of them in the antigenic supersite recognized by potent neutralizing mAbs, suggest that the NTD is under selective pressure from the host humoral immune response. This is further supported by the identification of NTD deletions within the antigenic supersite in immunocompromised hosts with prolonged infections (Avanzato et al., 2020; Choi et al., 2020; McCarthy et al., 2020), the *in vitro* selection of SARS-CoV-2 S escape variants with NTD mutations that decrease the neutralization potency of COVID-19 convalescent patient sera (Andreano et al., 2020; Weisblum et al., 2020) and the escape mutants reported here. We propose that mAb cocktails comprising S2L28 plus S2M28, S2×28, or S2×333 could provide broad protection against SARS-CoV-2 variants, such as the B.1.1.7 and 501Y.V2 lineages, as these mAbs are susceptible to distinct sets of SARS-CoV-2 S mutations, as previously proposed for RBD-specific mAbs (Greaney et al., 2020).

Here, we show that site i-targeting NTD neutralizing mAbs efficiently activate FcγRIIa and FcγRIIIa *in vitro*. Fc-mediated effector functions can be affected by the epitope specificity of the mAbs (Piccoli et al., 2020), highlighting the importance of the orientation of the S-bound Fc fragments for efficient FcγR cross-linking and engagement. Presumably, due to the different angle of approach, the site vi-targeting NTD mAb S2M24 did not activate either FcγRIIa or FcγRIIIa. The contribution of Fc-mediated effector functions could further enhance the prophylactic activity of potent NTD-specific mAbs against SARS-CoV-2 in humans as demonstrated for two RBD-specific SARS-CoV-2 neutralizing mAbs (Schäfer et al., 2021; Winkler et al., 2020).

Our cryoEM structures of SARS-CoV-2 S bound to the ultrapotent RBD-specific mAb S2M11 and to S2×333, S2M28, or S2L28, together with evidence of additive neutralizing effect of RBD- and NTD-targeted mAbs, provide proof-of-concept for implementing countermeasures using mAb cocktails targeting both the NTD and the RBD and potentially also for vaccine design. As several examples of single amino acid mutations reducing or completely abrogating neutralization by immune sera have been reported (Li et al., 2020; Liu et al., 2020b; Weisblum et al., 2020), we propose that combinations of mAbs targeting distinct domains will reduce the likelihood of emergence of escape mutants, as previously demonstrated for MERS-CoV (Wang et al., 2018). Our work suggests that the potency and resistance to antigenic drift of RBD-focused subunit vaccines (Walls et al., 2020a) might benefit from implementing a multi-pronged approach focusing the immune response on both the RBD and the NTD (Zhang et al., 2020).

## Acknowledgements

We thank Hideki Tani (University of Toyama) for providing the reagents necessary for preparing VSV pseudotyped viruses. The authors would also like to thank Roberto Spreafico (GSK, Belgium) for initial SARS-CoV-2 NTD conservation analysis and Marcel Meury for assistance with protein production. This study was supported by the National Institute of General Medical Sciences (R01GM120553 to D.V.), the National Institute of Allergy and Infectious Diseases (DP1AI158186 and HHSN272201700059C to D.V.), a Pew Biomedical Scholars Award (D.V.), Investigators in the Pathogenesis of Infectious Disease Awards from the Burroughs Wellcome Fund (D.V.), Fast Grants (D.V.), the Natural Sciences and Engineering Research Council of Canada (M.M.), the University of Washington Arnold and Mabel Beckman cryoEM center and beamline 5.0.1 at the Advanced Light Source at Lawrence Berkley National Laboratory.

## Author Contributions

Experiment Design: M.M., A.D.M., F.L., D.C., M.S.P., and D.V.; Donors’ Recruitment and Sample Collection: E.C., F.B., and A.R.; Sample Processing: A.D.M., D.P., M.B., F.Z., and C.S.F.; Protein expression and purification: M.M., SKZ, JEB, E.C., G.S., and D.V.; Isolation and Characterization of mAbs: M.M., A.D.M., F.L., M.A.T, D.P., A.C.W., M.B., A.C., Z.L., F.Z., S.Z., J.D.I., J.E.B., M.M.R., J.Z., L.E.R., S.B., B.G., C.S.F., R.A., S.C.F., P.W.R, L.M.B., F.B., and E.C.; binding and neutralization assays: M.M. A.D.M., F.L., M.A.T, D.P., A.C.W., M.B., A.C., Z.L., F.Z., M.M.R.; J.Z.; BLI/SPR assays: A.D.M, and L.E.R.; Cryo-EM Data Collection, Processing, and Model Building: M.M. and D.V.; Bioinformatic analysis of virus variants: J.D.I, and A.T.; Escape mutants selection and sequencing: Z.L., P.W.R, L.M.B, and S.P.J.W.; hamster model and data analysis: R.A., S.C.F., F.B., J.N., D.C., and M.S.P.; Data analysis: M.M., A.D.M, F.L., D.C., M.S.P., and D.V; Manuscript Writing: M.M., A.D.M., A.T., G.S., S.P.J.W, H.W.V, D.C., M.S.P., and D.V.; Supervision: F.B., E.C., J.N., A.R., G.S., A.T., S.P.J.W, H.W.V, D.C., M.S.P., and D.V.; Funding Acquisition: D.V.

