## Supplementary material for "N-terminal domain antigenic mapping reveals a site of vulnerability for SARS-CoV-2": Methods

**Table S1. NTD-specific mAb summary table**

| **#** | **Donor** | **mAb** | **IgVH  gene** | **HCDR3 length** | **VH % GL** | **IgVL  gene** | **VL % GL** | **ELISA vs NTD  (EC50 ng/ml)** | **Antigenic site** | **KD  NTD (M)** | **NT IgG vs  MLV-S2 pp (IC50 ng/ml)** | **Mx % NT** | **NT IgG vs live virus  IC50 (ng/ml)** | **Mx % NT** | **NT Fab vs live virus IC50 (ng/ml)** | **Mx % NT** |
| --- | --- | --- | --- | --- | --- | --- | --- | --- | --- | --- | --- | --- | --- | --- | --- | --- |
| 1 | **L** | S2L11 | 2-70 | 21 | 98.63 | K2-28 | 100 | 148.7 | ii | 7.25E-09 | nn | nn |  |  |  |  |
| 2 |  | S2L12 | 3-21 | 19 | 98.26 | K3-15 | 97.49 | 8.943 | iii | na | nn | nn |  |  |  |  |
| 3 |  | S2L14 | 1-69 | 18 | 98.61 | L4-69 | 98.64 | 507.4 | iii | 1.18E-08 | nn | nn |  |  |  |  |
| 4 |  | S2L20 | 3-30 | 15 | 97.92 | K1-33 | 96.42 | 447.7 | iv | 3.05E-08 | 2982 | 98% |  |  |  |  |
| 5 |  | S2L24 | 1-24 | 14 | 98.96 | K1-27 | 99.28 | 602.3 | i | 1.17E-08 | 49.1 | 83.50% |  |  |  |  |
| 6 |  | S2L26 | 1-24 | 14 | 97.22 | L3-10 | 98.21 | 47.72 | i | 4.56E-09 | 19.41 | 77% |  |  |  |  |
| 7 |  | **S2L28** | 3-21 | 19 | 96.53 | L2-14 | 97.57 | 34.26 | i | 1.20E-07 | 76.38 | 77% | 26.2 | 98.20% | 928 | 55.60% |
| 8 |  | S2L35 | 4-30 | 21 | 96.9 | L1-51 | 98.25 | 22.1 | iii | 7.63E-09 | nn | nn |  |  |  |  |
| 9 |  | S2L38 | 3-23 | 17 | 97.22 | K3-11 | 97.13 | 26.06 | iii | 1.62E-09 | nn | nn |  |  |  |  |
| 10 |  | S2L50 | 4-59 | 20 | 95.44 | L3-25 | 98.92 | 63.72 | i | 5.96E-09 | 24.88 | 91% |  |  |  |  |
| 11 | **M** | S2M24 | 4-61 | 20 | 97.25 | K1-39 | 95.34 | 698.7 | vi | 6.98E-09 | nn | nn |  |  |  |  |
| 12 |  | **S2M28** | 3-33 | 12 | 97.57 | L3-25 | 97.85 | 31.66 | i | 6.89E-09 | 53.73 | 88% | 5 | 98.50% | 26.3 | 66% |
| 13 | **X** | S2X15 | 1-3 | 21 | 98.98 | K3-11 | 96.14 | 12.3 | iii | 1.87E-09 | nn | nn |  |  |  |  |
| 14 |  | **S2X28** | 3-30 | 18 | 97.92 | L3-10 | 99.64 | 337.6 | i | na | 90 | 91% | 9.1 | 99.10% | 248.9 | 87.30% |
| 15 |  | S2X49 | 2-26 | 14 | 98.66 | L1-40 | 98.98 | 18.5 | iii | 4.97E-09 | nn | nn |  |  |  |  |
| 16 |  | S2X51 | 3-21 | 20 | 98.95 | K3-15 | 98.29 | 14.2 | iii | 3.24E-08 | nn | nn |  |  |  |  |
| 17 |  | S2X72 | 3-21 | 21 | 97.21 | K3-15 | 98.93 | 19.4 | iii | 2.19E-08 | nn | nn |  |  |  |  |
| 18 |  | S2X90 | 4-4 | 13 | 97.61 | L2-23 | 96.94 | 11.5 | iii | 3.35E-09 | nn | nn |  |  |  |  |
| 19 |  | S2X91 | 1-24 | 21 | 97.28 | K2-24 | 97.64 | 60.5 | i | 3.39E-09 | 40.3 | 87% |  |  |  |  |
| 20 |  | S2X93 | 3-33 | 20 | 97.97 | K3-15 | 96.51 | 13.4 | iii | 6.71E-08 | nn | nn |  |  |  |  |
| 21 |  | S2X94 | 3-53 | 14 | 97.6 | K3-15 | 98.59 | 8.7 | iii | 3.73E-08 | nn | nn |  |  |  |  |
| 22 |  | S2X98 | 3-21 | 24 | 97.61 | K3-15 | 99.65 | 20 | iii | 6.14E-08 | nn | nn |  |  |  |  |
| 23 |  | S2X102 | 4-59 | 15 | 97.81 | K3-20 | 96.81 | 24.3 | iii | 8.23E-09 | nn | nn |  |  |  |  |
| 24 |  | S2X105 | 4-59 | 17 | 98.29 | K3-15 | 99.3 | 12.4 | iii | 1.39E-08 | nn | nn |  |  |  |  |
| 25 |  | S2X107 | 4-38 | 16 | 96.95 | K1-39 | 96.83 | 69 | i | 4.08E-08 | 7.9 | 65% |  |  |  |  |
| 26 |  | S2X124 | 3-30 | 22 | 98.98 | K1-13 | 98.25 | 30 | i | 4.70E-08 | 8.9 | 68% |  |  |  |  |
| 27 |  | S2X125 | 3-21 | 17 | 95.6 | L1-40 | 97.57 | 24 | iii | 3.39E-09 | nn | nn |  |  |  |  |
| 28 |  | S2X158 | 1-24 | 16 | 96.25 | L1-47 | 95.91 | 18.7 | i | 8.54E-09 | 32.2 | 96% |  |  |  |  |
| 29 |  | S2X161 | 1-46 | 21 | 95.46 | L3-25 | 99.64 | 16.4 | i | 8.42E-09 | 17 | 83% |  |  |  |  |
| 30 |  | S2X165 | 4-61 | 20 | 96.65 | L10-54 | 96.59 | 51.8 | i | 3.46E-08 | 61.07 | 98% |  |  |  |  |
| 31 |  | S2X169 | 3-23 | 17 | 97.96 | L1-51 | 98.64 | 10.9 | iii | 1.49E-08 | nn | nn |  |  |  |  |
| 32 |  | S2X170 | 3-21 | 20 | 96.96 | K3-15 | 97.9 | 26.7 | iii | 1.62E-08 | nn | nn |  |  |  |  |
| 33 |  | S2X173 | 4-59 | 15 | 97.95 | L3-21 | 97.21 | 52.8 | v | 1.22E-08 | nn | nn |  |  |  |  |
| 34 |  | S2X175 | 3-21 | 20 | 97.72 | K3-15 | 96.167 | 16.8 | iii | 8.94E-09 | nn | nn |  |  |  |  |
| 35 |  | S2X176 | 2-70 | 10 | 98.65 | K4-1 | 98.01 | 474.6 | ii | 1.08E-08 | nn | nn |  |  |  |  |
| 36 |  | S2X186 | 1-2 | 12 | 98.97 | L2-23 | 98.62 | 18.3 | iii | 9.58E-09 | nn | nn |  |  |  |  |
| 37 |  | S2X303 | 2-5 | 17 | 95.88 | L3-1 | 95.34 | 14.2 | i | 6.30E-09 | 79.1 | 85% |  |  |  |  |
| 38 |  | S2X310 | 4-34 | 20 | 97.14 | L2-23 | 96.53 | 8 | iii | 5.58E-09 | nn | nn |  |  |  |  |
| 39 |  | S2X316 | 3-48 | 18 | 99.66 | K2-29 | 94.73 | 97.8 | v | 1.77E-08 | nn | nn |  |  |  |  |
| 40 |  | S2X320 | 3-33 | 17 | 96.53 | K1-33 | 97.85 | 26.9 | i | 1.81E-08 | 147.9 | 42% |  |  |  |  |
| 41 |  | **S2X333** | 3-33 | 17 | 96.53 | L3-21 | 97.49 | 7.6 | i | 2.89E-08 | 43.3 | 94% | 3 | 98.70% | 6.1 | 82.20% |

**Table S2. Cryo-EM data collection, refinement and validation statistics**

|  | S + S2M11 + **S2X333** | | S + S2M11 + **S2M28** | | S + S2M11 + **S2L28** | |
| --- | --- | --- | --- | --- | --- | --- |
| **Data collection and processing** |  | |  | |  | |
| Magnification (nominal) | 130,000 | | 130,000 | | 130,000 | |
| Voltage (kV) | 300 | | 300 | | 300 | |
| Electron exposure (e^–^/Å^2^) | 70 | | 70 | | 70 | |
| Defocus range (μm) | 0.8-2.0 | | 0.8-2.0 | | 0.8-2.0 | |
| Pixel size (Å) | 0.896 | | 1.05 | | 1.05 | |
| Processing Type | Global | Local | Global | Local | Global | Local |
| Symmetry imposed | C3 | C1 | C3 | C1 | C3 | C1 |
| Initial particle images (no.) | 490,229 | 785,859 | 383,144 | 665,979 | 356,514 | 473,310 |
| Final particle images (no.) | 261,953 | 99,627 | 221,993 | 282,884 | 157,770 | 81,887 |
| Map resolution (Å)  FSC threshold | 2.2  0.143 | 2.8  0.143 | 2.5  0.143 | 2.6  0.143 | 2.6  0.143 | 3.0  0.143 |
| **Refinement** |  |  |  |  |  |  |
| Initial model used (PDB code) | 7K43 | 6ZGE | 7K43 | 6ZGE | 7K43 | 6ZGE |
| Model resolution (Å) |  |  |  | 2.6 |  | 3.0 |
| FSC threshold | 0.143 | 0.143 | 0.143 | 0.143 | 0.143 | 0.143 |
| Map sharpening *B* factor (Å^2^) | -56 | -35 | -69 | -26 | -70 | -23 |
| Model composition |  |  |  |  |  |  |
| Nonhydrogen atoms | 33,615 | 2,596 | 34,041 | 3,384 | 34,641 | 3,281 |
| Protein residues | 4,470 | 321 | 4,563 | 414 | 4,590 | 409 |
| Glycan residues | 75 | 6 | 72 | 8 | 78 | 7 |
| *B* factors (Å^2^) |  |  |  |  |  |  |
| Protein | 24 | 24 | 20 | 99 | 23 | 30 |
| Glycans | 29 | 32 | 22 | 132 | 28 | 37 |
| R.m.s. deviations |  |  |  |  |  |  |
| Bond lengths (Å) | 1.05 | 1.00 | 1.02 | 1.02 | 1.02 | 0.96 |
| Bond angles (°) | 0.01 | 0.01 | 0.01 | 0.01 | 0.01 | 0.01 |
| **Validation** |  |  |  |  |  |  |
| MolProbity score | 0.71 | 0.50 | 0.68 | 0.66 | 0.81 | 0.84 |
| Clashscore | 0.62 | 0.00 | 0.52 | 0.45 | 0.8 | 0.47 |
| Rotamer outliers (%) | 0.09 | 0.00 | 0.09 | 0.56 | 0.00 | 0.00 |
| Ramachandran plot |  |  |  |  |  |  |
| Favored (%) | 98 | 98 | 98 | 99 | 98 | 97 |
| Allowed (%) | 2 | 2 | 2 | 1 | 2 | 3 |
| EMRinger score | 5.2 | 4.8 | 4.7 | 4.1 | 4.6 | 4.3 |
| **Data Availability** |  |  |  |  |  |  |
| EMDB |  |  |  |  |  |  |
| PDB |  |  |  |  |  |  |

**Table S3. Data collection and refinement statistics**

|  | **SARS-CoV-2 NTD + S2M28 Fab** |
| --- | --- |
| **Data collection** |  |
| Facility | ALS |
| Beamline | 5.0.2 |
| Wavelength (Å) | 0.97741 |
| Space group | *P* 2_1_ 2_1_ 2_1_ |
| Cell dimensions |  |
| *a*, *b*, *c* (Å) | 66.4 125.5 365.9 |
| *ɑ, β, ɣ* (°) | 90, 90, 90 |
| Resolution Range (Å) | 50-3.0 (3.1-3.0) |
| *R*_Sym_ *^†^* | 0.19 (2.16) |
| *R*_pim_ *^†^* | 0.05 (0.60) |
| *CC* ^†^* | 1.00 (0.91) |
| *I / σI* | 12.60 (1.39) |
| Completeness (%) | 100 (100) |
| Redundancy | 13.3 (13.5) |
| **Refinement** |  |
| No. reflections | 62431 |
| *R*_work_ / *R*_free_ *^‡^* | 0.223 / 0.239 |
| No. atoms |  |
| Protein | 11009 |
| Ligand/ion | 465 |
| Water | 0 |
| *B*-factors |  |
| Protein | 92 |
| Ligand/ion | 142 |
| R.m.s. deviations |  |
| Bond lengths (Å) | 1.00 |
| Bond angles (°) | 0.005 |
| Ramachandran Statistics |  |
| Favored | 98 |
| Allowed | 100 |

Note: Values in parentheses correspond to the highest resolution shell.

*^†^*, *R*_Sym_ =∑∑| *I* – (*I*) | /∑∑*I*, *R*_Pim_ =∑√(1/(n-1)∑| *I* – (*I*) | /∑∑*I*, and CC* =√(2CC_1/2_/(1+CC_1/2_)) where CC_1/2_ is the Pearson correlation coefficient of two half data sets as described elsewhere (Karplus and Diederichs, 2012)

*^‡^, R*_work_ = ∑ | |*F*_obs_| - k|*F*_calc_| | / |*F*_obs_| where *F*_obs_ and *F*_calc_ are the observed and calculated structure factors, respectively. *R*_free_ is the sum extended over a subset of reflections (5%) excluded from all stages of the refinement.

**Table S4. Additional cryo-EM data collection, refinement and validation statistics**

|  | 2P-DS-S +  **S2X28** | S + S2M11 +  **S2L20** | S + S2M11 +  **S2X316** | S + S2M11 +  **S2M24** |
| --- | --- | --- | --- | --- |
| **Data collection** |  |  |  |  |
| Magnification (nominal) | 130,000 | 36,000 | 130,000 | 130,000 |
| Voltage (kV) | 300 | 200 | 200 | 200 |
| Electron exposure (e^–^/Å^2^) | 70 | 60 | 60 | 60 |
| Defocus range (μm) | 0.8-2.0 | 0.8-2.0 | 0.8-2.0 | 0.8-2.0 |
| Pixel size (Å) | 1.05 | 1.16 | 1.16 | 1.16 |
| **Processing** |  |  |  |  |
| Symmetry imposed | C1 | C3 | C3 | C3 |
| Initial particle images (no.) | 140,562 | 8345 | 11,695 | 5840 |
| Final particle images (no.) | 36,413 | 3588 | 4,340 | 3,770 |
| Map resolution (Å)  FSC threshold | ~4-5  0.143 | 6.3  0.143 | 6.0  0.143 | 6.5  0.143 |
| Map sharpening *B* factor (Å^2^) | -28 | -264 | -168 | -255 |
| **Data Availability** |  |  |  |  |
| EMDB |  |  |  |  |

**Key Resources Table**

| **REAGENT or RESOURCE** | **SOURCE** | **IDENTIFIER** |
| --- | --- | --- |
| **Bacterial Strains** | | |
| *E. coli* DH10B Competent Cells | ThermoFisher Scientific | Cat# EC0113 |
| **Deposited Data** | | |
| S + S2M11 + S2X333 CryoEM Map | https://www.ebi.ac.uk/pdbe/emdb/ |  |
| S + S2M11 + S2X333 CryoEM Model | https://www.rcsb.org |  |
| S + S2M11 + S2X333 Focused CryoEM Map | https://www.ebi.ac.uk/pdbe/emdb/ |  |
| S + S2M11 + S2X333 Focused CryoEM Model | https://www.rcsb.org |  |
| S + S2M11 + S2M28 CryoEM Map | https://www.ebi.ac.uk/pdbe/emdb/ |  |
| S + S2M11 + S2M28 CryoEM Model | https://www.rcsb.org |  |
| S + S2M11 + S2M28 Focused CryoEM Map | https://www.ebi.ac.uk/pdbe/emdb/ |  |
| S + S2M11 + S2M28 Focused CryoEM Model | https://www.rcsb.org |  |
| S + S2M11 + S2L28 CryoEM Map | https://www.ebi.ac.uk/pdbe/emdb/ |  |
| S + S2M11 + S2L28 CryoEM Model | https://www.rcsb.org |  |
| S + S2M11 + S2L28 Focused CryoEM Map | https://www.ebi.ac.uk/pdbe/emdb/ |  |
| S + S2M11 + S2L28 Focused CryoEM Model | https://www.rcsb.org |  |
| 2P-DS-S + S2X28 CryoEM Map | https://www.ebi.ac.uk/pdbe/emdb/ |  |
| S + S2M11 + S2L20 CryoEM Map | https://www.ebi.ac.uk/pdbe/emdb/ |  |
| S + S2M11 + S2X316 CryoEM Map | https://www.ebi.ac.uk/pdbe/emdb/ |  |
| S + S2M11 + S2M24 CryoEM Map | https://www.ebi.ac.uk/pdbe/emdb/ |  |
| S + S2M11 + S2X176 CryoEM Map | https://www.ebi.ac.uk/pdbe/emdb/ |  |
| S + S2M11 + S2X94 CryoEM Map | https://www.ebi.ac.uk/pdbe/emdb/ |  |
| S NTD + S2M28 Crystal Structure | https://www.rcsb.org |  |
| **Experimental Models: Cell Lines** | | |
| FreeStyle™ 293-F Cells | ThermoFisher Scientific | Cat# R79007 |
| Expi293F™ Cells | ThermoFisher Scientific | Cat# A14527 |
| HEK293T | ATCC | Cat# CRL-11268 |
| Vero-E6 | ATCC | CRL-1586 |
| Expi-CHO cells | ThermoFisher Scientific | Cat# A29127 |
| Expi-CHO cells stably expressing SARS-CoV-2 | This study | N/A |
| Jurkat cells stably expressing FcγRIIIa receptor (V158 variant) and NFAT-driven luciferase gene (effector cells) | Promega | Cat# G7018 |
| Jurkat cells stably expressing FcγRIIa receptor (H131 variant) and NFAT-driven luciferase gene | Promega | Cat# G9995 |
| **Oligonucleotides** | | |
| NTD_fwd: TATAATGGTACCGCCACCATGGGCAT | Integrated DNA Technologies, Inc. | N/A |
| NTD_rev: TATAATAAGCTTTCAGTGGTGGTGGTGGTGGTGATGATGCGCGGTAAAGCTCTTCGCTGTACACTT | Integrated DNA Technologies, Inc. | N/A |
| N149Q_fwd: AAAAGTCCTGGATGGAGTCTGAGT | Integrated DNA Technologies, Inc. | N/A |
| N149Q_rev: GGTTCTTGTGATAGTACACGCCC | Integrated DNA Technologies, Inc. | N/A |
| D253G_fwd: GCTCCTCTAGCGGATGGAC | Integrated DNA Technologies, Inc. | N/A |
| D253G_rev: CGCCTGGTGTCAGGTAGG | Integrated DNA Technologies, Inc. | N/A |
| D253Y_fwd: ACTCCTCTAGCGGATGGAC | Integrated DNA Technologies, Inc. | N/A |
| D253Y_rev: AGCCTGGTGTCAGGTAGG | Integrated DNA Technologies, Inc. | N/A |
| T19A_fwd: ACAAGGACCCAGCTGCC | Integrated DNA Technologies, Inc. | N/A |
| T19A_rev: GGCCAGGTTCACGCACTG | Integrated DNA Technologies, Inc. | N/A |
| R246A_fwd: CATCCTACCTGACACCAGGC | Integrated DNA Technologies, Inc. | N/A |
| R246A_rev: CGTGCAGGGCCAGCAG | Integrated DNA Technologies, Inc. | N/A |
| L18F_fwd: CTTTACCACAAGGACCCAGC | Integrated DNA Technologies, Inc. | N/A |
| L18F_rev: TTCACGCACTGTGTGCC | Integrated DNA Technologies, Inc. | N/A |
| H146Y_fwd: ACAAGAACAATAAGTCCTGGATGGAGTC | Integrated DNA Technologies, Inc. | N/A |
| H146Y_rev: AATAGTACACGCCCAGGAATGGAT | Integrated DNA Technologies, Inc. | N/A |
| A222V_fwd: TCCTGGAGCCACTGGTG | Integrated DNA Technologies, Inc. | N/A |
| A222V_rev: CGGAGAATCCCTGTGGCAG | Integrated DNA Technologies, Inc. | N/A |
| SP_rev: AGGACGAGGAAGACGAACATGGTGGCGGTACCTCCG | Integrated DNA Technologies, Inc. | N/A |
| SP_fwd: GCTCCCTCTGGTTTCTTCCCAGTGCGTGAACCTGACCA | Integrated DNA Technologies, Inc. | N/A |
| SP_S12P_fwd: GCTCCCTCTGGTTCCATCCCAGTGCGTGAACCTGACCA | Integrated DNA Technologies, Inc. | N/A |
| Y144del_fwd: CACAAGAACAATAAGTCCTGGATGGAGT | Integrated DNA Technologies, Inc. | N/A |
| Y144del_rev: GTACACGCCCAGGAATGGATC | Integrated DNA Technologies, Inc. | N/A |
| S254F_fwd: CTCTAGCGGATGGACCGCA | Integrated DNA Technologies, Inc. | N/A |
| S254F_rev: AAGTCGCCTGGTGTCAGGTAG | Integrated DNA Technologies, Inc. | N/A |
| K147T_fwd: CGAACAATAAGTCCTGGATGGAGTCT | Integrated DNA Technologies, Inc. | N/A |
| K147T_rev: TGTGATAGTACACGCCCAGGAAT | Integrated DNA Technologies, Inc. | N/A |
| C136Y_fwd: ATAATGATCCATTCCTGGGCGTGTA | Integrated DNA Technologies, Inc. | N/A |
| C136Y_rev: AAAACTGGAACTCGCACACCTTG | Integrated DNA Technologies, Inc. | N/A |
| **Recombinant DNA** | | |
| pCMV::SARS-CoV-2_S_ecto_hexapro | Ref: PMID: 32577660 | N/A |
| pCMV::SARS-CoV-2_S_ecto_2P_DS | Ref PMID: 32753755 | N/A |
| pCMV::SARS-CoV-2_S_NTD | This study | N/A |
| pCMV::P-GD_S_ecto | This study | N/A |
| pCMV::SARS-CoV-2_S_ecto_avi | PMID: 32972994 | N/A |
| pcDNA3.1::SARS-CoV-2-S-D19 | (Ou et al., 2020) | N/A |
| phCMV1::SARS-CoV-2 | PMID: 32972994 | N/A |
| phCMV1::RatG13_S | This study | N/A |
| phCMV1::P-GX_S | This study | N/A |
| phCMV1::P-GD_S | This study | N/A |
| phCMV1::ZC45_S | This study | N/A |
| phCMV1::ZXC21_S | This study | N/A |
| phCMV1::YN2013_S | This study | N/A |
| phCMV1::SARS-CoV_S | This study? | N/A |
| phCMV1::RmYN02_S | This study | N/A |
| phCMV1::Bt-kY72_S | This study | N/A |
| phCMV1::BM48-31_S | This study | N/A |
| pSARS-CoV-2-Nluc | (Xie et al., 2020) |  |
| **Commercial products** |  |  |
| PEI MAX | Polysciences | Cat# POL24765-1 |
| 4-Nitrophenyl phosphate disodium salt hexahydrate (pNPP) | Sigma-Aldrich | Cat# N2765-100TAB |
| Tween 20 | Sigma-Aldrich | Cat# 93773 |
| Bovine Serum Albumine (BSA) | Sigma-Aldrich | Cat# 3059 |
| TPCK treated Trypsin | Worthington Biochem | Cat# LS003750 |
| Alexa Fluor 647-AffiniPure F(ab’)2  Fragment Goat Anti-Human IgG, Fcg  Fragment Specific | Jackson ImmunoResearch | Cat# 109-606-098 |
| Goat Anti-Human IgG-AP | Southern Biotech | Cat# 2040-04 |
| Streptavidin, Alexa Fluor 647 conjugate | Life technologies | Cat# S21374 |
| Bio-Glo | Promega | Cat# G7940 |
| **Software and Algorithms** | | |
| cryoSPARC v3.0.1 | (Punjani et al., 2017) | https://cryosparc.com |
| Relion v3.0 | (Zivanov et al., 2018) | https://www3.mrc-lmb.cam.ac.uk/relion |
| Coot | (Casanal et al., 2019) | https://www2.mrc-lmb.cam.ac.uk/personal/pemsley/coot/ |
| Phenix-Refine | (Adams et al., 2010) | https://www.phenix-online.org/download/ |
| Phenix-Phaser | (McCoy et al., 2007) | https://www.phenix-online.org/download/ |
| XDS | (Kabsch, 2010) | http://xds.mpimf-heidelberg.mpg.de |
| Prism 8 | GraphPad Software | https://www.graphpad.com/scientific-software/prism/ |
| Biacore Evaluation software | Cytiva | https://www.cytivalifesciences.com/en/us/shop/protein-analysis/spr-label-free-analysis/software/biacore-insight-evaluation-software-p-23528 |

**Cell lines**

Cell lines used in this study were obtained from ATCC (HEK293T and Vero-E6)or ThermoFisher Scientific (Expi CHO cells, FreeStyle™ 293-F cells and Expi293F™ cells).

**Sample donors**

Samples were obtained from three SARS-CoV-2 recovered individuals (L, M and X) under study protocols approved by the local Institutional Review Boards (Canton Ticino Ethics Committee, Switzerland, the Ethical committee of Luigi Sacco Hospital, Milan, Italy). All donors provided written informed consent for the use of blood and blood components (such as PBMCs, sera or plasma).

Samples were collected 14 and 52 days after symptoms onset for donor L and M, respectively. Blood drawn from donor X was obtained at day 36, 48, 75 and 125 after symptoms onset.

**Cloning and mutant generation**

SARS-CoV-2 NTD was sub-cloned with E. coli DH10B Competent Cells into pCMV using primers NTD_fwd and NTD_rev. The resulting construct was mutated by PCR mutagenesis to generate N149Q, D253G/Y, T19A, R246A, L18F, H146Y, A222V, Y144del, S254F, K147T, C136Y, and the NTD construct with native signal peptide with and without S12P, using the eponymously named primers (**Key Resources Table**). The genes encoding for the Sarbecovirus S proteins tested were cloned in the phCMV1 or pcDNA.3 vectors, and the gene for the C-terminally his-tagged ectodomain of P-GD S was cloned into pCMV (**Key Resources Table**). Plasmid sequences were verified by Genewiz sequencing facilities (Brooks Life Sciences).

**Recombinant ectodomains production**

All SARS-CoV-2 S spike ectodomains were produced in 500 mL cultures of FreeStyle™ 293-F cells (ThermoFisher Scientific) grown in suspension using FreeStyle 293 expression medium (ThermoFisher Scientific) at 37°C in a humidified 8% CO2 incubator rotating at 130 r.p.m. Cells grown to a density of 2.5 million cells per mL were transfected using PEI (9 μg/mL) and pCMV::SARS-CoV-2_S_ecto_hexapro, pCMV::SARS-CoV-2_S_ecto_2P_DS, pCMV::P-GD_S_ecto, pCMV::SARS-CoV-2_S_ecto_avi, pCMV::SARS-CoV-2_S_D614G_ecto_avi and cultivated for 4 days. The supernatant was harvested and cells were resuspended for another three days, yielding two harvests. S ectodomains were purified from clarified supernatants using a Cobalt affinity column (Cytiva, HiTrap TALON crude), washing with 20 column volumes of 20 mM Tris-HCl pH 8.0 and 150 mM NaCl and eluted with a gradient of 600 mM imidazole. The same protocol was followed for P-GD spike ectodomain purification, except that 25 mM sodium phosphate pH 7 and 300 mM sodium chloride were used instead of 20 mM Tris-HCl pH 8.0 and 150 mM NaCl. At this stage, SARS-CoV-2 S with the avi tag (from pCMV::SARS-CoV-2_S_ecto_avi) was biotinylated (BirA biotin-protein ligase standard reaction kit, Avidity) and further purified by size exclusion chromatography (Superose6, GE Healthcare). All purified proteins were then concentrated using a 100 kDa centrifugal filter (Amicon Ultra 0.5 mL centrifugal filters, MilliporeSigma), residual imidazole was washed away by consecutive dilutions in the centrifugal filter unit with 20 mM Tris-HCl pH 8.0 and 150 mM NaCl, and finally concentrated to 5 mg/ml and flash frozen.

All SARS-CoV-2 S NTD domain constructs (residues 14-307) with a C-terminal 8XHis-tag were produced in 100 mL culture of Expi293F™ Cells (ThermoFisher Scientific) grown in suspension using Expi293™ Expression Medium (ThermoFisher Scientific) at 37°C in a humidified 8% CO2 incubator rotating at 130 r.p.m.) (Walls et al., 2020) (Walls et al., 2020) (Walls et al., 2020) (Walls et al., 2020) (Walls et al., 2020) (Walls et al., 2020) (Walls et al., 2020) (Walls et al., 2020) (Walls et al., 2020) (Walls et al., 2020) (Walls et al., 2020). Cells grown to a density of 3 million cells per mL were transfected using pCMV::SARS-CoV-2_S_NTD derivative mutants with the ExpiFectamine™ 293 Transfection Kit (ThermoFisher Scientific) with and cultivated for five days at which point the supernatant was harvested. His-tagged NTD domain constructs were purified from clarified supernatants using 2 ml of cobalt resin (Takara Bio TALON), washing with 50 column volumes of 20 mM HEPES-HCl pH 8.0 and 150 mM NaCl and eluted with 600 mM imidazole. Purified protein was concentrated using a 30 kDa centrifugal filter (Amicon Ultra 0.5 mL centrifugal filters, MilliporeSigma), the imidazole was washed away by consecutive dilutions in the centrifugal filter unit with 20 mM HEPES-HCl pH 8.0 and 150 mM NaCl, and finally concentrated to 20 mg/ml and flash frozen. For crystallization, the purified NTD was not frozen but was further purified by size exclusion chromatography (Superdex Increase 75 10/300 G, GE Healthcare), concentrated using a new 30 kDa centrifugal filter, and used immediately.

**Intact mass spectrometry analysis of purified NTD constructs**

The purpose of intact MS was to verify the n-terminal sequence on four constructs. N-linked glycans were removed by PNGase F after overnight non-denaturing reaction at room temperature. 4μg of deglycosylated protein was used for each injection on the LC-MS system to acquire intact MS signal after separation of protease and protein by LC (Agilent PLRP-S reversed phase column). Thermo MS (Q Exactive Plus Orbitrap) was used to acquire intact protein mass under denaturing condition. BioPharma Finder 3.2 software was used to deconvolute the raw m/z data to protein average mass.

**Non-reducing Peptide Mapping mass spectrometry analysis of purified NTD constructs**

The purpose of peptide mapping was to identify the post modification on unpaired cysteines. Tryptic digestion was used without adding reducing reagent. 50μg of deglycosyated protein was denatured (6M guanidine hydrochloride), alkylated (Iodoacetamide), and buffer exchanged (Zeba spin desalting column) before trypsin digestion. 10ug of digested peptide was analyzed on the LC-MS system (Agilent AdvanceBio peptide mapping column and Thermo Q Exactive Plus Orbitrap MS) to acquire both MS1 and MS2 data under HCD fragmentation. Peptide mapping data was analyzed on Biopharma Finder 3.2 by searching the possible modifications such as cysteinylation and carbamidomethylation on the cysteines.

**Isolation of peripheral blood mononuclear cells (PBMCs), plasma and sera**

PBMCs were isolated from blood draw performed using tubes pre-filled with heparin, followed by Ficoll density gradient centrifugation. PBMCs were either used freshly along SARS-CoV2 Spike protein specific memory B cells sorting or stored in liquid nitrogen for later use. Sera were obtained from blood collected using tubes containing clot activator, followed by centrifugation and stored at -80 °C.

**B-cell isolation and recombinant mAb production**

Starting from freshly isolated PBMCs or upon cells thawing, B cells were enriched by staining with CD19 PE-Cy7 (BD Bioscience 341113) and incubation with anti-PE bead (Miltenyi Biotec, cat. 130- 048-801), followed by positive selection using LS columns. Enriched B cells were stained with anti-IgM, anti-IgD, anti-CD14 and anti-IgA, all PE labelled, and prefusion SARS-CoV-2 S with a biotinylated avi tag conjugated to Streptavidin Alexa-Fluor 647 (Life Technologies). SARSCoV-2 S-specific IgG+ memory B cells were sorted by flow cytometry via gating for PE negative and Alexa-Fluor 647 positive cells. Cells were cultured for the screening of positive supernatants. Antibody VH and VL sequences were obtained by RT-PCR and mAbs were expressed as recombinant human Fab fragment or as IgG1 (G1m3 allotype) carrying the half-life extending M428L/N434S (LS) mutation in the Fc region. ExpiCHO cells were transiently transfected with heavy and light chain expression vectors as previously described (Pinto et al., 2020).

Affinity purification was performed on ÄKTA Xpress FPLC (Cytiva) operated by UNICORN software version 5.11 (Build 407) using HiTrap Protein A columns (Cytiva) for full length human and hamster mAbs and CaptureSelect CH1-XL MiniChrom columns (ThermoFisher Scientific) for Fab fragments, using PBS as mobile phase. Buffer exchange to the appropriate formulation buffer was performed with a HiTrap Fast desalting column (Cytiva). The final products were sterilized by filtration through 0.22 µm filters and stored at 4 ºC.

**Enzyme-linked immunosorbent assay (ELISA)**

To determine specificity of recombinantly produced mAbs, 96 half area well-plates (Corning) were coated over-night at 4°C with of SARS-CoV-2 S, NTD or RBD proteins prepared 1 μg/ml, 2 μg/ml and 5 μg/ml in PBS pH 7.2, respectively. Plates were then blocked with PBS 1% BSA (Sigma) and subsequently incubated with mAbs serial dilutions for 1 h at room temperature. After 2 washing steps with PBS 0.05% Tween 20 (PBS-T) (Sigma-Aldrich) goat anti-huma IgG secondary antibody (Southern Biotech) was added in incubated for 1 h at room temperature. Plates were then washed again with PBS-T and 4-NitroPhenyl phosphate (pNPP, Sigma-Aldrich) substrate added. After 30 min incubation, absorbance at 405 nm was measured by a plate reader (Biotek) and data plotted using Prism GraphPad.

For all other applications reported, the following ELISA procedure was followed: 30 µl of ectodomains (stabilized prefusion trimer) of S or NTD from SARS-CoV-2 were coated on 384 well ELISA plates at 1 ng/µl for 16 hours at 4°C. Plates were washed with a 405 TS Microplate Washer (BioTek Instruments) then blocked with 80 µl SuperBlock (PBS) Blocking Buffer (Thermo Scientific) for 1 hour at 37°C. Plates were then washed and 30 µl antibodies were added to the plates at concentrations between 0.001 and 100,000 ng/ml and incubated for 1 h at 37°C. Plates were washed and then incubated with 30 µl of 1/5000 diluted goat anti-human Fc IgG-HRP (invitrogen A18817). Plates were washed and then 30 µl Substrate TMB microwell peroxidase (Seracare 5120-0083) was added for 4 min at room temperature. The colorimetric reaction was stopped by addition of 30 µl of 1 N HCl. A_450_ was read on a Varioskan Lux plate reader (Thermo Scientific).

**MLV-based pseudovirus production and neutralization**

To generate SARS-CoV-2 S murine leukemia virus pseudotyped virus, HEK293T cells were seeded in 10-cm dishes in DMEM supplemented with 10% FBS. The next day cells were transfected with a SARS-CoV-2 S glycoprotein-encoding plasmid harboring the D19 C-terminal truncation (Ou et al., 2020) using the X-tremeGENE HP DNA transfection reagent (Roche) according to the manufacturer’s instructions. Cells were then incubated at 37°C with 5% CO_2_ for 72 h. Supernatant was harvested and cleared from cellular debris by centrifugation at 400 X g, and stored at -80 °C.

For neutralization assays, Vero E6 cells were seeded into white 96-well plates (PerkinElmer) at 20,000 cells/well and cultured overnight at 37 °C with 5 % CO_2_ in 100 µl DMEM supplemented with 10% FBS and 1% penicillin/streptomycin. The next day, MLV-SARS-CoV-2 pseudovirus was activated with 10 μg/ml TPCK treated-Trypsin (Worthington Biochem) for 1 h at 37 °C. Then recombinant antibodies at various concentrations were incubated with activated pseudovirus for 1 h at 37 °C. The Vero E6 cells were then washed with DMEM, and the 50 μl of pseudovirus/mAbs mixes were added and incubated for 2 h at 37 °C with 5 % CO_2_. After incubation, 50 µl of DMEM containing 20% FBS and 2 % penicillin/streptomycin was added and the cells were incubated 48 h at 37 °C with 5 % CO_2_. Following these 48 h of infection, culture medium was removed from the cells and 50 µl/wellof Bio-Glo (Promega) (diluted 1:2 with PBS with Ca2+Mg2+ (Thermo Fisher) was added to the cells and incubated in the dark for 15 min before reading on a Synergy H1 Hybrid Multi-Mode plate reader (Biotek). Measurements were done in duplicate and RLU values were converted to percentage of neutralization and plotted with a nonlinear regression curve fit in Graph Prism.

**Neutralization of authentic SARS-CoV-2-Nluc virus**

Neutralization of authentic SARS-CoV-2 by entry-inhibition assay Neutralization was determined using SARS-CoV-2-Nluc, an infectious clone of SARSCoV-2 (based on strain 2019-nCoV/USA_WA1/2020) which encodes nanoluciferase in place of the viral ORF7 and demonstrated comparable growth kinetics to wildtype virus (Xie et al., 2020). Vero E6 cells were seeded into black-walled, clear-bottom 96-well plates at 2 x 104 cells/well and cultured overnight at 37 ºC. The next day, 9-point 4-fold serial dilutions of mAbs were prepared in infection media (DMEM + 10% FBS). SARS-CoV-2-Nluc was diluted in infection media at a final MOI of 0.1 or 0.01 PFU/cell, added to the mAb dilutions and incubated for 30 minutes at 37 ºC. Media was removed from the Vero E6 cells, mAb-virus complexes were added and incubated at 37 ºC for 6 or 24 hours. Media was removed from the cells, Nano-Glo luciferase substrate (Promega) was added according to the manufacturer’s recommendations, incubated for 10 minutes at room temperature and the luciferase signal was quantified on a VICTOR Nivo plate reader (Perkin Elmer).

**Binding and affinity determination by Biolayer Interferometry (BLI)**

BLI measurements were performed using an Octet Red96 (ForteBio). All reagents were prepared in kinetics buffer (PBS plus 0.01% BSA) at the indicated concentrations.

BLI was used to assess antibody binding affinity to SARS-CoV-2 NTD. IgG antibodies were prepared at 2.7 μg/ml and captured on pre-hydrated Protein A biosensors (Sartorius) for 1 min. The biosensors with immobilized antibodies were moved into kinetics buffer with SARS-CoV-2 NTD (concentrations tested: 333.3, 166.6, 83.3, 41.7, 20.8, 10.4, 5.2 nM) for 5 min (i.e. association). The dissociation of the SARS-CoV-2 NTD was then recorded for 9 min in wells containing kinetics buffer. Affinity constants were calculated using a global fit model and results were plotted using GraphPad Prism.

BLI was also used to assess antibody competition studies to define the NTD antigenic map. Biotinylated SARS-CoV-2 S protein was prepared at 10 µg/ml in kinetics buffer and loaded on pre-hydrated High Precision Streptavidin SAX Biosensors (Sartorius) for 3 min. NTD mAbs at 20 µg/ml in kinetics buffer were then sequentially added to observe binding competition and signal recorded for 5 min (or 7 min)

BLI was also used to assess mAb-mediated inhibition of SARS-CoV-2 S binding to human recombinant ACE2. Before the assay SARS-CoV2 S ectodomain trimer (5 µg/ml) was incubated with tested mAbs (30 µg/ml) or no mAb for 30 minutes at 37°C. Biotinylated recombinant human ACE2 protein (2 µg/ml) was immobilized on High Precision Streptavidin SAX Biosensors (Sartorius). Next, an association step with S/mAb complexes was performed for 10 minutes. Results were plotted using GraphPad Prism.

**Affinity determination by Surface Plasmon Resonance (SPR)**

SPR binding measurements were performed using a Biacore T200 instrument where purified avi-tagged SARS-CoV-2 S D614G ectodomain trimer was captured using anti-AviTag pAb covalently immobilized on a CM5 sensor chip. The running buffer was Cytiva HBS-EP+ pH 7.4; measurements were performed at 25˚C. Affinity/avidity determinations were run as single-cycle kinetics, with a 3-fold dilution series of mAb starting from 300 nM, and each concentration injected for 180 sec. Double reference-subtracted data were fit to a 1:1 binding model using Biacore Evaluation software. Fit results for IgG yielded apparent equilibrium dissociation constants due to avidity. For dissociation rates that were too slow to fit, equilibrium dissociation constants are reported as an upper limit.

**Transient Expression of Sarbecovirus S protein in ExpiCHO-S Cells.**

Immediately before transfection, ExpiCHO-S cells were seeded at 6 x 10^6^ cells cells/mL in a volume of 5 mL in a 50 mL bioreactor. Spike coding plasmids were diluted in cold OptiPRO SFM, mixed with ExpiFectamine CHO Reagent (Life Technologies) and added to the cells. Transfected cells were then incubated at 37°C with 8% CO_2_ with an orbital shaking speed of 120 RPM (orbital diameter of 25 mm) for 42 hours

**Binding to cell surface expressed Sarbecovirus S proteins by Flow Cytometry**

Transiently transfected ExpiCHO cells were harvested and washed two times in wash buffer (PBS 1% BSA, 2 mM EDTA). Cells were counted and distributed into round bottom 96-well plates (Corning) and incubated with the NTD antibodies at the final concentration of 5 μg/ml. Alexa Fluor647-labelled Goat Anti-Human IgG secondary Ab (Jackson Immunoresearch) was prepared at 1.5 μg/ml added onto cells after two washing steps. Cells were then washed twice and resuspended in wash buffer for data acquisition at ZE5 cytometer (Biorad).

**Fusion inhibition assay**

Vero E6 cells were seeded in 96 well plates at 15,000 cells per well in 70 μl DMEM with high glucose and 2.4% FBS (Hyclone). After 16 h at 37 °C with 8 % CO_2_, the cells were transfected with SARS-CoV-2-S-D19_pcDNA3.1 as follows: for 10 wells, 0.57 µg plasmid SARS-CoV-2- S-D19_pcDNA3.1 were mixed with 1.68 µl X-tremeGENE HP in 30 µl OPTIMEM. After 15 minutes incubation, the mixture was diluted 1:10 in DMEM medium and 30μl was added per well. A 4-fold serial dilution mAbs was prepared and added to the cells, with a starting concentration of 20 μg/ml. The following day, 30 μl 5X concentrated DRAQ5 in DMEM was added per well and incubated for 2 hours at 37˚C. Nine images of each well were acquired with a Cytation 5 equipment for analysis.

**Measurement of Fc-effector functions**

mAb-dependent activation of human FcγRIIIa was performed with a bioluminescent reporter assay. ExpiCHO cells stably expressing full-length wild-type SARS-CoV-2 S (target cells) were incubated with different amounts of mAbs. After a 15-minute incubation, Jurkat cells stably expressing FcγRIIIa receptor (V158 variant) or FcγRIIa receptor (H131 variant) and NFAT-driven luciferase gene (effector cells) were added at an effector to target ratio of 6:1 for FcγRIIIa and 5:1 for FcγRIIa. Signaling was quantified by the luciferase signal produced as a result of NFAT pathway activation. Luminescence was measured after 20 hours of incubation at 37˚C with 5% CO2 with a luminometer using the Bio-Glo-TM Luciferase Assay Reagent according to the manufacturer’s instructions (Promega, Cat. Nr.: G9798, G7018 and G9995).

**Cell-surface mAb-mediated S_1_ shedding**

CHO cells stably expressing wild-type SARS-CoV-2 S were resuspended in wash buffer (PBS 1 % BSA, 2 mM EDTA) and treated with 10 μg/mL TPCK-trypsin (Worthington Biochem) for 30 min at 37°C. Cells were then washed and distributed into round bottom 96-well plates (90,000 cells/well). MAbs were added to cells at 15 µg/mL final concentration for 180 min at 37 ºC. Cells were collected at different time points (5, 30, 60, 120 and 180), washed with wash buffer at 4 ºC, and incubated with 1.5 mg/mL secondary goat anti-human IgG, Fc fragment specific (Jackson ImmunoResearch) on ice for 20 min. Cells were washed and resuspended in wash buffer and analyzed with ZE5 FACS (Bio-rad).

**Cryo-EM sample preparation and data collection**.

2.5 µL of 12 mg/mL S2M11 Fab, 2.5 µL of 12 mg/mL NTD Fab, and 6 µL of 5 mg/mL SARS-CoV-2 S (specifically the hexapro construct (Hsieh et al., 2020)) were incubated together for 30 min at 37 ºC. Alternatively, 3 µL of 3 mg/mL S2X28 Fab, and 3 µL of 1 mg/mL SARS-CoV-2 2P DS S (McCallum et al., 2020) were incubated together for 30 min at 37 ºC. Unbound Fab was then washed away with three consecutive dilutions in 400 µL of 20 mM Tris-HCl pH 8.0 and 150 mM NaCl over a 100 kDa centrifugal filter (Amicon Ultra 0.5 mL centrifugal filters, MilliporeSigma). The complex was concentrated to 1.2 mg/ml and 3 µL was applied onto a freshly glow discharged 2.0/2.0 UltraFoil grid (200 mesh), plunge frozen using a vitrobot MarkIV (ThermoFisher Scientific) using a blot force of -1 and 6.5 second blot time at 100% humidity and 23°C.

Data were acquired using the Leginon software(Suloway et al., 2005) to control a FEI Titan Krios or Glacios transmission electron microscope equipped with a Gatan K2 Summit direct detectors and operated at 300 kV with a Gatan Quantum GIF energy filter or at 200 kV, respectively. For both microscopes, the dose rate was adjusted to 8 counts/pixel/s, and each movie was acquired in 50 frames of 200 ms. For the Krios, ~3000 micrographs were acquired for each session with a super-resolution pixel size of 0.525 Å with a defocus range between -0.8 and -2.0 μm. For the Glacios, ~100 micrographs with a pixel size of 1.16 Å were collected in a single session with a defocus range between -0.8 and -2.0 μm.

**Cryo-EM data processing**

Movie frame alignment, estimation of the microscope contrast-transfer function parameters, particle picking and extraction (with a box size of 400 pixels^2^) were carried out using Warp(Tegunov and Cramer, 2019). At this stage, the pixel size was binned to 1.05 Å for data collected from the Krios. Reference-free 2D classification was performed using cryoSPARC(Punjani et al., 2017) to select well-defined particle images. 3D classification with 50 iterations each (angular sampling 7.5˚ for 25 iterations and 1.8˚ with local search for 25 iterations) were carried out using Relion(Zivanov et al., 2018) without imposing symmetry to separate distinct SARS-CoV-2 S conformations. 3D refinements were carried out using non-uniform refinement along with per-particle defocus refinement in cryoSPARC(Punjani et al., 2017). Particle images were subjected to Bayesian polishing(Zivanov et al., 2019). To accommodate the resolution of the S/S2M11/S2X333 data collection, at the polishing stage the box size was adjusted to 600 Å while the pixel size was binned to 0.896 Å. Another round of non-uniform refinement in cryoSPARC(Punjani et al., 2017) was performed, followed by global and per-particle defocus refinement and again non-uniform refinement. Reported resolutions are based on the gold-standard Fourier shell correlation (FSC) of 0.143 criterion and Fourier shell correlation curves were corrected for the effects of soft masking by high-resolution noise substitution(Scheres and Chen, 2012).

**Cryo-EM model building and analysis**

UCSF Chimera(Goddard et al., 2007) and Coot (Casanal et al., 2019) were used to fit atomic models (PDB 7K43 and 6ZGE) into the cryo-EM maps. The model was then refined into the map using Rosetta(DiMaio et al., 2015; Frenz et al., 2019; Wang et al., 2016) and analyzed using MolProbity(Chen et al., 2010), EMringer(Barad et al., 2015), and Phenix(Liebschner et al., 2019). Figures were generated using UCSF ChimeraX(Goddard et al., 2018) and UCSF Chimera(Goddard et al., 2007).

**S2M28 Fab and NTD co-crystallization and structure determination**

Crystals of S2M28 Fab with SARS-CoV-2 NTD were identified by the sitting-drop vapor diffusion method, set-up by hand with MCSG-1 crystallization screen (Anatrace). Optimized crystals were generated with 1 µl of 5.7 mg/ml Fab and 4 mg/ml NTD in 20 mM HEPES-HCl pH 8.0 and 150 mM NaCl plus 1 µl mother liquor solution containing 0.2 M Ammonium Sulfate, 0.1 M Sodium Citrate pH 4.75, 25 % (w/v) PEG4000. Crystals were flash cooled in liquid nitrogen using the mother liquor solution supplemented with 17.5 % (w/v) xylitol as a cryoprotectant. Diffraction data were collected on synchrotron beamline 5.0.2 at the Advanced Light Source, and processed with the XDS software package (Kabsch, 2010). Initial phases were obtained by molecular replacement using Phenix-Phaser (McCoy et al., 2007), using the S/S2M11/S2M28 cryoEM structure. Several subsequent rounds of model building and refinement were performed using Coot (Casanal et al., 2019) and Phenix-Refine (Adams et al., 2010).

**Data availability**

The cryoEM maps, crystallographic data, and atomic models will be deposited to the EMDB and PDB.

Adams, P.D., Afonine, P.V., Bunkoczi, G., Chen, V.B., Davis, I.W., Echols, N., Headd, J.J., Hung, L.W., Kapral, G.J., Grosse-Kunstleve, R.W.*, et al.* (2010). PHENIX: a comprehensive Python-based system for macromolecular structure solution. Acta Crystallogr D Biol Crystallogr *66*, 213-221.

Barad, B.A., Echols, N., Wang, R.Y., Cheng, Y., DiMaio, F., Adams, P.D., and Fraser, J.S. (2015). EMRinger: side chain-directed model and map validation for 3D cryo-electron microscopy. Nat Methods *12*, 943-946.

Casanal, A., Lohkamp, B., and Emsley, P. (2019). Current Developments in Coot for Macromolecular Model Building of Electron Cryo-microscopy and Crystallographic Data. Protein Sci.

Chen, V.B., Arendall, W.B., 3rd, Headd, J.J., Keedy, D.A., Immormino, R.M., Kapral, G.J., Murray, L.W., Richardson, J.S., and Richardson, D.C. (2010). MolProbity: all-atom structure validation for macromolecular crystallography. Acta Crystallogr D Biol Crystallogr *66*, 12-21.

DiMaio, F., Song, Y., Li, X., Brunner, M.J., Xu, C., Conticello, V., Egelman, E., Marlovits, T., Cheng, Y., and Baker, D. (2015). Atomic-accuracy models from 4.5-Å cryo-electron microscopy data with density-guided iterative local refinement. Nat Methods *12*, 361-365.

Frenz, B., Rämisch, S., Borst, A.J., Walls, A.C., Adolf-Bryfogle, J., Schief, W.R., Veesler, D., and DiMaio, F. (2019). Automatically Fixing Errors in Glycoprotein Structures with Rosetta. Structure *27*, 134-139.e133.

Goddard, T.D., Huang, C.C., and Ferrin, T.E. (2007). Visualizing density maps with UCSF Chimera. Journal of structural biology *157*, 281-287.

Goddard, T.D., Huang, C.C., Meng, E.C., Pettersen, E.F., Couch, G.S., Morris, J.H., and Ferrin, T.E. (2018). UCSF ChimeraX: Meeting modern challenges in visualization and analysis. Protein Sci *27*, 14-25.

Hsieh, C.L., Goldsmith, J.A., Schaub, J.M., DiVenere, A.M., Kuo, H.C., Javanmardi, K., Le, K.C., Wrapp, D., Lee, A.G., Liu, Y.*, et al.* (2020). Structure-based design of prefusion-stabilized SARS-CoV-2 spikes. Science.

Kabsch, W. (2010). XDS. Acta Crystallogr D Biol Crystallogr *66*, 125-132.

Karplus, P.A., and Diederichs, K. (2012). Linking crystallographic model and data quality. Science *336*, 1030-1033.

Liebschner, D., Afonine, P.V., Baker, M.L., Bunkóczi, G., Chen, V.B., Croll, T.I., Hintze, B., Hung, L.W., Jain, S., McCoy, A.J.*, et al.* (2019). Macromolecular structure determination using X-rays, neutrons and electrons: recent developments in Phenix. Acta Crystallogr D Struct Biol *75*, 861-877.

McCallum, M., Walls, A.C., Bowen, J.E., Corti, D., and Veesler, D. (2020). Structure-guided covalent stabilization of coronavirus spike glycoprotein trimers in the closed conformation. Nat Struct Mol Biol.

McCoy, A.J., Grosse-Kunstleve, R.W., Adams, P.D., Winn, M.D., Storoni, L.C., and Read, R.J. (2007). Phaser crystallographic software. J Appl Crystallogr *40*, 658-674.

Pinto, D., Park, Y.J., Beltramello, M., Walls, A.C., Tortorici, M.A., Bianchi, S., Jaconi, S., Culap, K., Zatta, F., De Marco, A.*, et al.* (2020). Cross-neutralization of SARS-CoV-2 by a human monoclonal SARS-CoV antibody. Nature *583*, 290-295.

Punjani, A., Rubinstein, J.L., Fleet, D.J., and Brubaker, M.A. (2017). cryoSPARC: algorithms for rapid unsupervised cryo-EM structure determination. Nat Methods *14*, 290-296.

Scheres, S.H., and Chen, S. (2012). Prevention of overfitting in cryo-EM structure determination. Nature methods *9*, 853-854.

Suloway, C., Pulokas, J., Fellmann, D., Cheng, A., Guerra, F., Quispe, J., Stagg, S., Potter, C.S., and Carragher, B. (2005). Automated molecular microscopy: the new Leginon system. Journal of structural biology *151*, 41-60.

Tegunov, D., and Cramer, P. (2019). Real-time cryo-electron microscopy data preprocessing with Warp. Nature methods.

Walls, A.C., Park, Y.J., Tortorici, M.A., Wall, A., McGuire, A.T., and Veesler, D. (2020). Structure, Function, and Antigenicity of the SARS-CoV-2 Spike Glycoprotein. Cell *181*, 281-292.e286.

Wang, R.Y., Song, Y., Barad, B.A., Cheng, Y., Fraser, J.S., and DiMaio, F. (2016). Automated structure refinement of macromolecular assemblies from cryo-EM maps using Rosetta. Elife *5*.

Xie, X., Muruato, A.E., Zhang, X., Lokugamage, K.G., Fontes-Garfias, C.R., Zou, J., Liu, J., Ren, P., Balakrishnan, M., Cihlar, T.*, et al.* (2020). A nanoluciferase SARS-CoV-2 for rapid neutralization testing and screening of anti-infective drugs for COVID-19. bioRxiv, 2020.2006.2022.165712.

Zivanov, J., Nakane, T., Forsberg, B.O., Kimanius, D., Hagen, W.J., Lindahl, E., and Scheres, S.H. (2018). New tools for automated high-resolution cryo-EM structure determination in RELION-3. Elife *7*.

Zivanov, J., Nakane, T., and Scheres, S.H.W. (2019). A Bayesian approach to beam-induced motion correction in cryo-EM single-particle analysis. IUCrJ *6*.
